# A Sequential Niche Multimodal Conformation Sampling Algorithm for Protein Structure Prediction

**DOI:** 10.1101/2020.12.29.424663

**Authors:** Yu-Hao Xia, Chun-Xiang Peng, Xiao-Gen Zhou, Gui-Jun Zhang

## Abstract

**Motivation:** Massive local minima on the protein energy surface often causes traditional conformation sampling algorithms to be easily trapped in local basin regions, because they are difficult to stride over high-energy barriers. Also, the lowest energy conformation may not correspond to the native structure due to the inaccuracy of energy models. This study investigates whether these two problems can be alleviated by a sequential niche technique without loss of accuracy.

**Results:** A sequential niche multimodal conformation sampling algorithm for protein structure prediction (SNfold) is proposed in this study. In SNfold, a derating function is designed based on the knowledge learned from the previous sampling and used to construct a series of sampling-guided energy functions. These functions then help the sampling algorithm stride over high-energy barriers and avoid the re-sampling of the explored regions. In inaccurate protein energy models, the high-energy conformation that may correspond to the native structure can be sampled with successively updated sampling-guided energy functions. The proposed SNfold is tested on 300 benchmark proteins and 24 CASP13 FM targets. Results show that SNfold is comparable with Rosetta restrained by distance (Rosetta-dist) and C-QUARK. SNfold correctly folds (TM-score ≥ 0.5) 231 out of 300 proteins. In particular, compared with Rosetta-dist protocol, SNfold achieves higher average TM-score and improves the sampling efficiency by more than 100 times. On the 24 CASP13 FM targets, SNfold is also comparable with four state-of-the-art methods in the CASP13 server group. As a plugin conformation sampling algorithm, SNfold can be extended to other protein structure prediction methods.

**Availability:** The source code and executable versions are freely available at https://github.com/iobio-zjut/SNfold.

## 1 Introduction

The stunning diversity of molecular functions performed by proteins is made possible by their finely tuned three-dimensional structures (Kuhlman and Bradley, 2019). *De novo* protein structure prediction is challenging because it requires both an accurate energetic representation of a protein structure and an efficient conformation sampling algorithm (Lee *et al.*, 2011), where the latter is the primary bottleneck that restricts the accuracy of *de novo* protein structure prediction (Bradley *et al.*, 2005; Dill and Maccallum, 2012; Kandathil *et al.*, 2016). Recently, geometric optimization (Senior *et al.*, 2020) has achieved remarkable success in this community, but Monte Carlo (MC) (Li and Scheraga, 1987; Hansmann and Okamoto, 1999) based fragment assembly is still an important method as it reveals the dynamic process of protein folding to some extent. Historically, MC is a popular algorithm for conformation sampling (Lee *et al.*, 2009), as well as its variants, such as Metropolis Monte Carlo (MMC) (Metropolis *et al.*, 1953; Kuhlman and Bradley, 2019), replica-exchange Monte Carlo (REMC) (Kihara *et al.*, 2001; Zhou *et al.*, 2019) and parallel hyperbolic sampling (PHS) (Zhang *et al.*, 2002). Rosetta (Rohl *et al.*, 2004; Park *et al.*, 2019), which involves MMC sampling based on fragment assembly in conjunction with knowledge-based energy functions, is one of the most popular suites for macromolecular modeling (Dukka, 2017). For a target sequence, Rosetta commonly requires executing a large number of independent MMC trajectories and using a cluster algorithm to identify the most frequently sampled conformations. QUARK (Xu and Zhang, 2012; Zheng *et al.*, 2019) is one of the top ranked servers in recent CASPs, in which models are assembled from fragments by REMC simulation under the guide of an atomic-level knowledge-based force field. However, the prediction accuracy of fragment assembly approaches has been observed to decrease for larger proteins (> 150 residues) because larger proteins require significantly more computing as the conformational space is vastly increased (Kim *et al.*, 2009; Moult *et al.*, 2018). In addition, MC algorithms are often trapped in local basin regions during conformation sampling. Therefore, MC algorithms based fragment assembly face a challenge on how to improve sampling efficiency without loss of accuracy.

Evolutionary algorithms (EAs) (Custodio *et al.*, 2014; Clausen and Shehu, 2015; Zhou *et al.*, 2019; Zhou and Zhang, 2019) are a class of powerful stochastic optimization algorithms, which have less of a chance of getting stuck at local minima compared to MC algorithms (Saleh *et al.*, 2013; Shehu, 2015). Some studies have been conducted for protein conformation sampling. MOEA (Olson and Shehu, 2013) combines local and global search in a population-based EA, and evolves a fixed-size population of conformations through a series of generations under the guidance of Pareto analysis. A hybridization protocol (Ovchinnikov *et al.*, 2018) is developed, in which the overall iterative process is guided by an EA applying hybridization as mutation or crossover operations and controlling diversity within the structural pool. In our recent research, CGLFold (Liu *et al.*, 2020) is proposed, in which a loop-specific local perturbation model is designed to improve the accuracy of models based on a differential evolution algorithm. However, population-based EAs inherently run the risk of premature convergence (Ovchinnikov *et al.*, 2018; Peng *et al.*, 2020). Fortunately, this problem can be alleviated with multimodal optimization strategies, and some related studies have been carried out. RMA (Garza-Fabre *et al.*, 2016) uses a stochastic ranking-based survival selection procedure to minimize the evaluation function while keeping the structural diversity. A multimodal memetic algorithm (Correa *et al.*, 2018) is proposed, and it involves an evolutionary approach with a ternary tree-structured population allied to a local search strategy. However, multimodal optimization is accompanied with expensive calculation costs. Meanwhile, sampling the native structure in cases of inaccurate energy models is difficult for these algorithms.

In recent years, significant progress has been witnessed on *de novo* protein structure prediction, which is mainly due to the success of sequence-based contact and distance predictions (Moult *et al.*, 2018; Kryshtafovych *et al.*, 2019). This alleviates the inaccuracy of energy models. In CASP13, DeepMind’s entry, AlphaFold (A7D), placed first in the free-modeling (FM) category (AlQuraishi, 2019). AlphaFold (Senior *et al.*, 2019, 2020) trains a neural network to accurately predict distances between pairs of residues, and then generates structures by using a stochastic gradient descent algorithm to optimize a potential constructed with distance information. RaptorX-Contact (Wang *et al.*, 2017; Xu, 2019; Xu and Wang, 2019) employs deep and fully convolutional residual neural network (ResNet) to predict distance distribution, secondary structure and backbone torsion angles. Then accurate models can be constructed quickly by feeding predicted restraints to crystallography and NMR system (CNS) (Brunger, 2007). CONFOLD2 (Adhikari and Cheng, 2018) utilizes predicted contacts and secondary structures to generate restraints that are used by the distance geometry and simulated annealing optimization implemented in CNS to build tertiary structure models. AmoebaContact (Mao *et al.*, 2020) adopts a set of network architectures optimized for contact prediction through automatic searching, and then GDFold (Mao *et al.*, 2020) considers all residue pairs from the prediction results of AmoebaContact in a differentiable loss function and optimizes atom coordinates by using the gradient descent algorithm to generate structures. Through the extension of deep learning-based prediction to inter-residue orientations in addition to distances, trRosetta (Yang *et al.*, 2020) supplements the predicted restraints with components of the Rosetta energy function to generate accurate models. These algorithms achieve a dramatical success by building a distance-based potential and then quickly constructing models via geometric optimization. Distance-assisted fragment assembly algorithms also show an excellent performance, which are still the mainstream, such as Rosetta (Park *et al.*, 2019) and C-QUARK (Zheng *et al.*, 2019). The reason may be that noisy contacts/distances can be offset by fragment assembly due to the fragments that come from known protein structures. However, sampling efficiency is rarely considered in distance-assisted fragment assembly algorithms in the existing literature. Therefore, it is a significant open problem that how to improve sampling efficiency while ensuring accuracy.

In this study, we propose a new conformation sampling algorithm, SNfold, which combines multimodal optimization with distance-assisted fragment assembly. Experimental results on the benchmark set with non-redundant proteins show that SNfold improves the sampling efficiency by more than 100 times compared with Rosetta-dist, and it is comparable with state-of-the-art methods in accuracy.

## 2 Methods

MC sampling and its variants have been extensively used in protein structure prediction (Park *et al.*, 2019; Zheng *et al.*, 2019). However, two broad problems should be considered: (1) the efficiency of conformation sampling; (2) the accuracy of energy function. For example, in MMC simulation (Figure 1), the first problem is that randomly starting multiple independent MMC trajectories may be inefficient. In Figure 1(a), both two MMC trajectories are trapped in the same local basin region, annotated as Basin 1, because of the high-energy barrier between Basin 1 and Basin 2, thereby causing redundant sampling in Basin 1. The second problem is the more serious case of the inaccurate energy function (Figure 1(b)), the conformation with the lowest energy is metastable, and the native structure is located in Basin 3. Therefore, even if the lowest energy region (Basin 2) is found, the native structure may not be sampled.

**Fig. 1.**
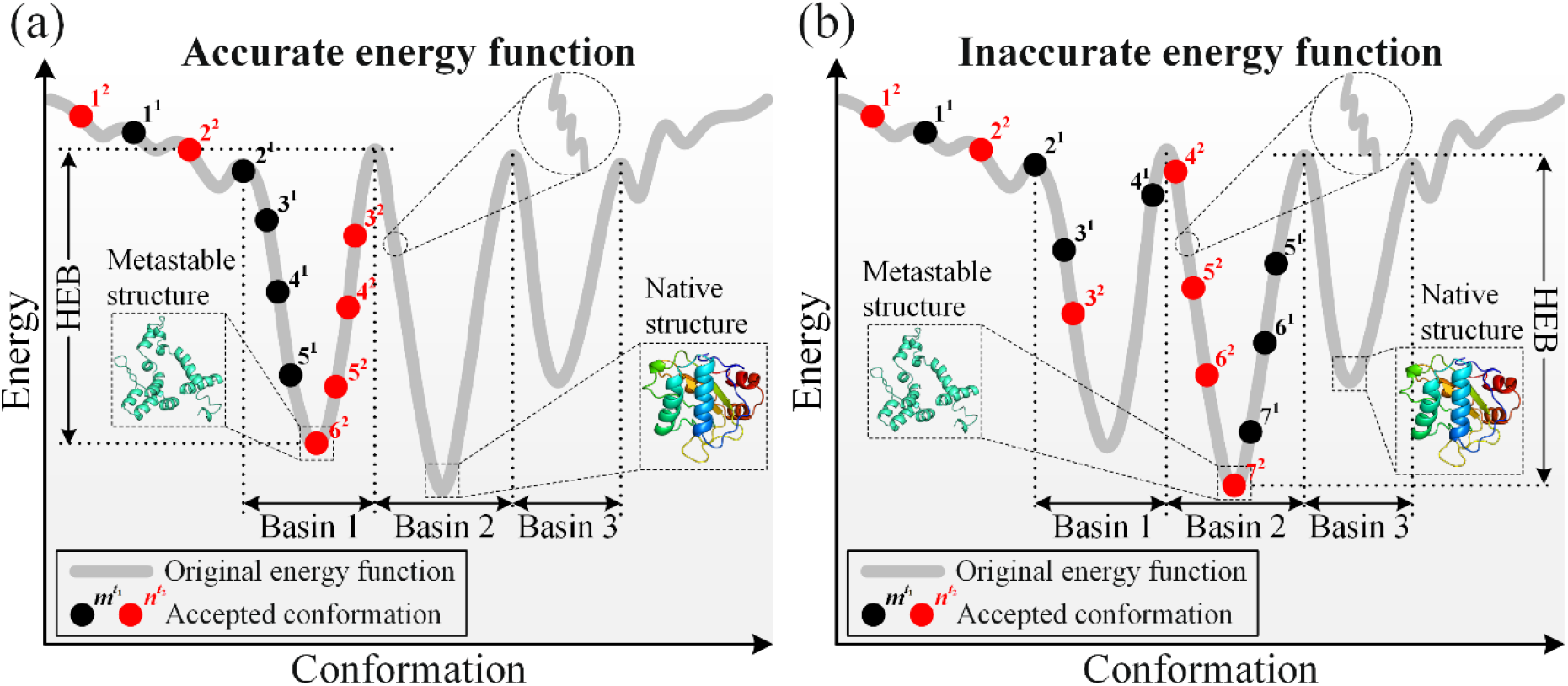
Schematic of two cases in MMC simulation. (a) Inefficient conformation sampling. (b) Inaccurate energy function. The black dot represents the *m*-th accepted conformation in the *t*_1_-th trajectory, and the red dot corresponds to the *n*-th accepted conformation in the *t*_2_ -th trajectory. Here, *t*_1_ = 1 and *t*_2_ = 1 are taken as an example. HEB stands for the high-energy barrier between basins, and the dotted circle refers to the details of the energy function, which is complicated and rugged.

For the two cases mentioned above, a sequential niche multimodal conformation sampling algorithm (SNfold) is proposed. In SNfold, a derating function is constructed on the basis of the knowledge learned from the previous sampling and applied to the original energy function to build a series of sampling-guided energy functions. With these functions, MMC simulation can easily stride over high-energy barriers. In terms of sampling efficiency, as shown in Figure 2(a), the second MMC trajectory strides over the barrier between Basin 1 and Basin 2 with the aid of the sampling-guided energy function, thereby avoiding the re-sampling of Basin 1. For the case of inaccurate energy function (Figure 2(b)), the native structure located in Basin 3 can be sampled more easily under the navigation of the sampling-guided energy function. Thus, the sampling efficiency of MMC simulation can be improved without loss of accuracy.

**Fig. 2.**
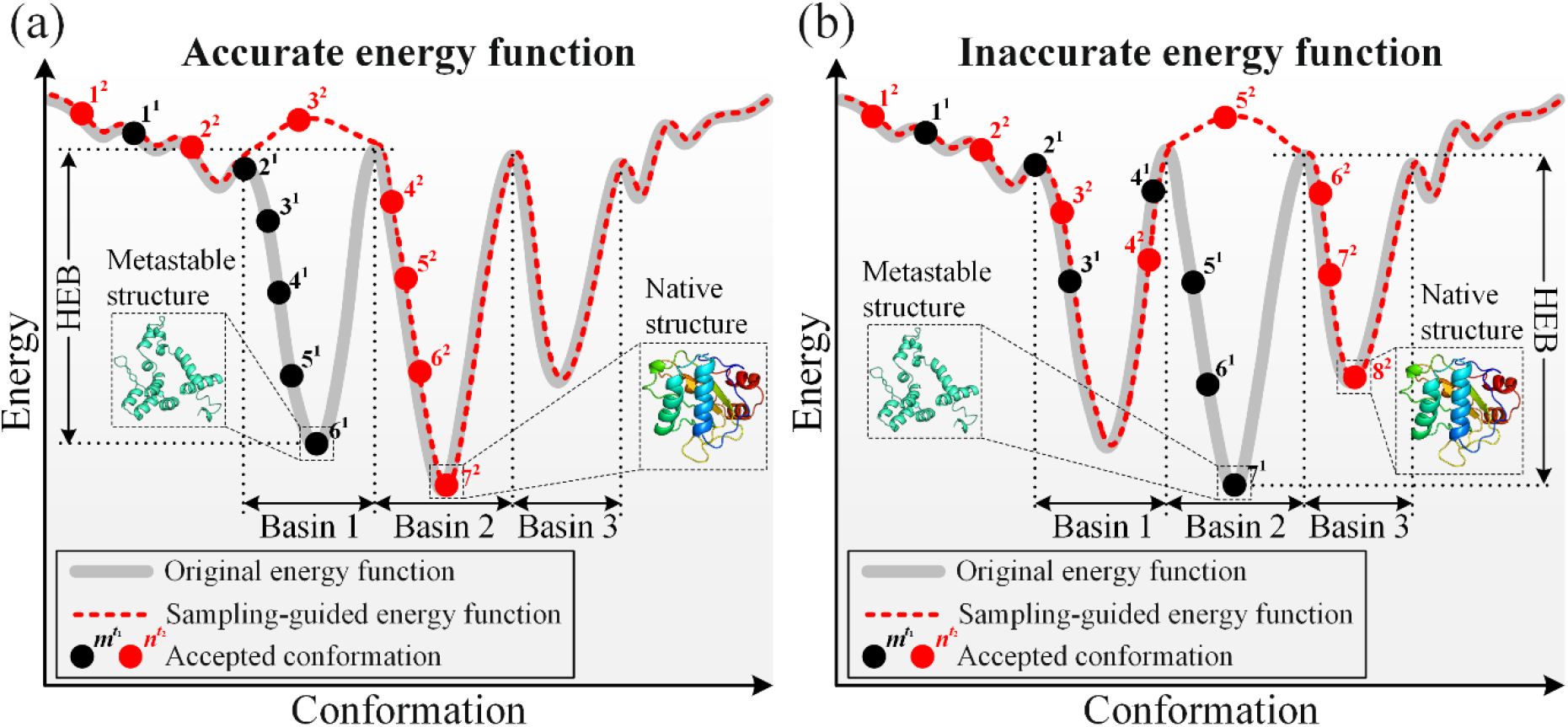
Schematic of MMC simulation in SNfold. (a) For the sampling efficiency, Basin 1 is filled under the action of the derating function, thereby avoiding the re-sampling of Basin 1 by the second MMC trajectory. (b) When the energy function is inaccurate, the native structure is sampled in the second MMC trajectory because Basin 2 is filled by the derating function.

The pipeline of SNfold is shown in Figure 3, and the flowchart of SNfold is presented in Supplementary Figure S1. In Figure 3, starting from a query sequence, the inter-residue distance distribution is predicted by the trRosetta server (https://yanglab.nankai.edu.cn/trRosetta/) and used to build the distance-based scoring function. Then, the initial conformations are generated by random fragment assembly, and the fragment library with homologues excluded is built by the Robetta fragment server (http://old.robetta.org/). Modal exploration contains the *T* MMC trajectories of Rosetta ClassicAbinitio protocol (Rohl *et al.*, 2004). For each MMC trajectory, the initial conformation is used as a starting point, and the conformation with the lowest energy in the trajectory is selected, called the seed conformation, *C*_seed_. A derating function is designed on the basis of the sampling knowledge (including seed conformation *C*_seed_ and niche radius *r*). The derating function is applied to the original energy function to construct a sampling-guided energy function, which is used to guide the next MMC simulation. In modal exploitation, the distance-based scoring function is designed to guide conformation sampling in basins where the seed conformations 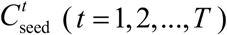 are located. Lastly, *T* models with the best distance score in the basin are selected.

**Fig. 3.**
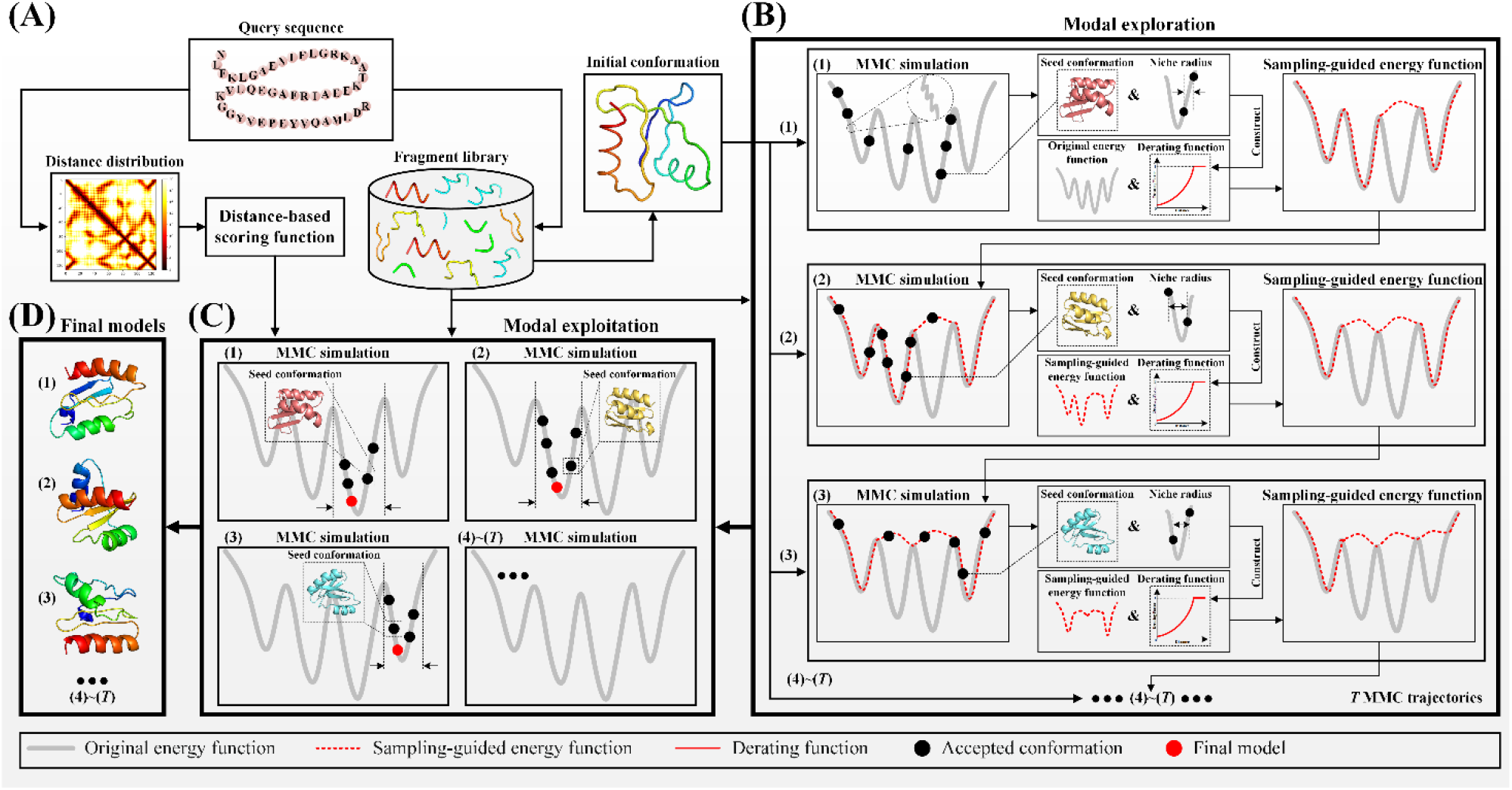
Pipeline of SNfold. (A) Conformation initialization. The initial conformations are generated by random fragment assembly, and the distance-based scoring function is constructed by the predicted distance by deep learning, which is used for modal exploitation. (B) Modal exploration. It consists of *T* MMC trajectories starting with the initial conformations. Based on the sampling knowledge (seed conformation and niche radius), a derating function is constructed to build the sampling-guided energy function, which is used to guide the next MMC simulation. (C) Modal exploitation. It contains *T* MMC trajectories with seed conformations as the initial, which are performed under the guidance of the original energy function and the distance-based scoring function. The sampling range is limited to the basin where the seed conformation is located. (D) Final prediction models.

According to the above strategy, a series of sampling-guided energy functions are generated without changing the original energy function. Consequently, MMC simulation can easily stride over high-energy barriers and avoid the redundant sampling of the explored basins. In addition, the likelihood of the native structure to be sampled is increased when the energy function is inaccurate.

### 2.1 Modal exploration

Searching for the lowest energy conformation may be unreliable because of the multimodality and inaccuracy of protein energy functions. Modal exploration is performed to navigate potential basins where the native structure may be located. Firstly, a similarity metric between two conformations is defined and used to determine the niche radius. Afterwards, a derating function is designed on the basis of the niche radius and seed conformation obtained from the previous sampling. Lastly, the derating function is utilized to construct a series of sampling-guided energy functions to guide subsequent sampling.

#### 2.1.1 Niche radius

A similarity metric is defined to describe the similarity between two conformations. Essentially, the similarity metric reflects the distance between two conformations in the conformational space. Given two conformations *C_m_* and *C_n_*, the similarity metric between them is defined as:

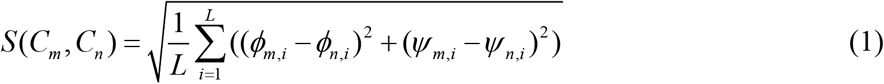

where *L* is the length of the protein sequence; *i* is the residue index; *ϕ_m, i_* and *ϕ_n, i_* are the dihedral angles about C-N-C*α*-C for the *i*-th residue of *C_m_* and *C_n_*, respectively; and *ψ_m, i_* and *ψ_n, t_* are the dihedral angles about N-C*α*-C-N for the *i*-th residue of *C_m_* and *C_n_*, respectively.

In each MMC trajectory, the basin where the seed conformation *C*_seed_ is located is determined as a region within the niche radius *r* centred on the seed conformation *C*_seed_ (Figure 4). The niche radius is calculated by:

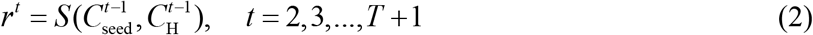

where 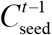 is the seed conformation of the (*t*-1)-th trajectory, and 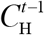 is the conformation with the highest energy at stages 3 and 4 of the Rosetta ClassicAbinitio protocol in the (*t*-1)-th trajectory. This selection is mainly attributed to our consideration that the conformations at stages 3 and 4 begin to converge towards the local basin (Rohl *et al.*, 2004; Garza-Fabre *et al.*, 2016). Therefore, the highest energy conformation at these two stages is considered to be at the edge of the basin where the seed conformation is located (Figure 4).

**Fig. 4.**
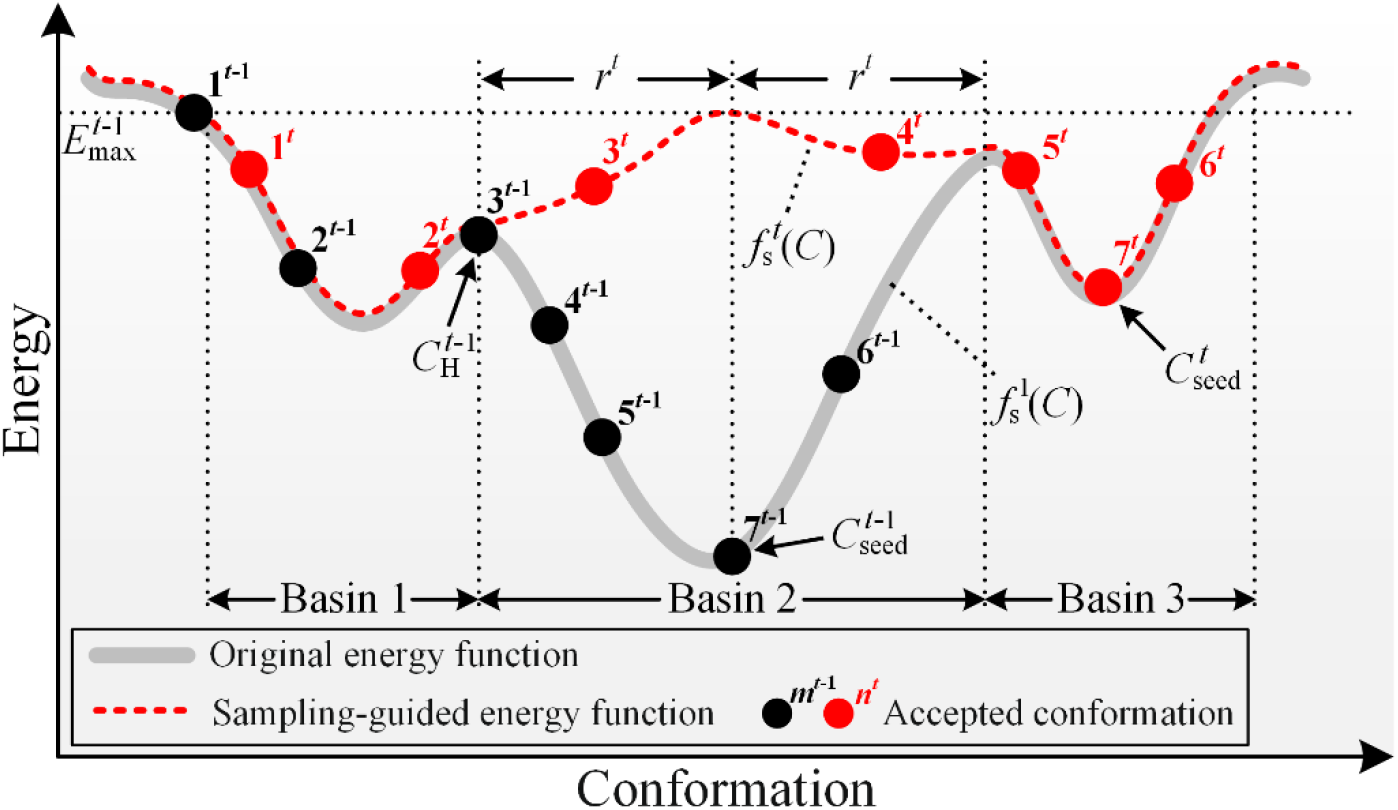
Schematic of the original energy function and the sampling-guided energy function. In the (*t*-1)-th MMC trajectory, 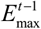 is equal to the energy value of the initial conformation because it has the highest energy among all the accepted conformations.

#### 2.1.2 Derating function

To avoid the re-sampling of previously explored basins in subsequent MMC trajectories, a derating function is designed based on the knowledge learned from the previous sampling (i.e., *C*_seed_ and *r*) in each trajectory. The derating function of the *t*-th trajectory is designed as:

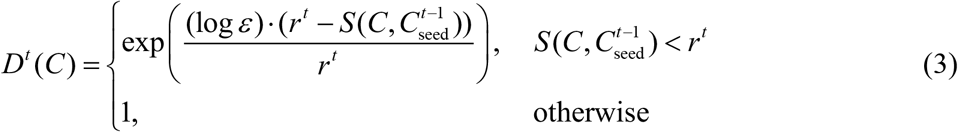

where *t* = 2,3,...,*T*; *C* is the target conformation; and 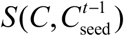 is the distance between the target conformation *C* and the seed conformation 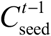, as determined by the similarity metric in Equation (1). The derating minimum value *ε*, is an arbitrarily small positive number, which determines the concavity of the derating curve, with smaller value of *ε* producing more concavity (Supplementary Figure S2). When 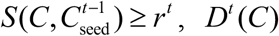 is set to 1.

#### 2.1.3 Sampling-guided energy function

By applying the derating function to the original energy function, a series of sampling-guided energy functions are constructed to guide the conformation sampling in subsequent trajectories. Here, the Rosetta score3 energy model (Rohl *et al.,* 2004) is selected as the original energy function and denoted as *E*_rosetta_. The sampling-guided energy function can be defined as:

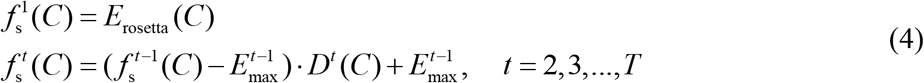

where *T* is the total number of MMC trajectories, 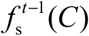 is the sampling-guided energy function of the (*t*-1)-th trajectory, and 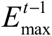. is a reference value, which is equal to the maximum energy value of all the accepted conformations in the (*t*-1)-th trajectory.

Essentially, the sampling-guided energy function is constructed by increasing the energy value of the previously explored basins in the original energy function by the derating function. The increase in energy value depends on the distances between the target conformation and the seed conformations of the previous trajectories. In Figure 4, 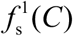 and 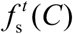 represent the original energy function and the sampling-guided energy function in the *t*-th trajectory, respectively. For 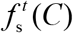, the basin explored in the (*t*-1)-th trajectory (Basin 2) is filled by the derating function *D^t^* (*C*), so that Basin 2 can be crossed more easily in the *t*-th trajectory. It is worth emphasizing that the derating function *D^t^* (*C*) does not destroy the shape of the original energy function 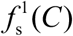, and 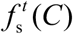 is used to guide conformation sampling in the *t*-th trajectory. If the distance between the target conformation *C* and 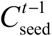 is less than 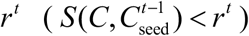, then *ε*≤ *D^t^* (*C*) < 1. According to Equation (4), 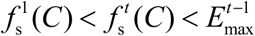. Therefore, in this case, the energy value of the target conformation *C* is raised, and the closer to 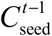, the higher the degree of increase. Specially, when 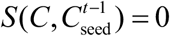, then 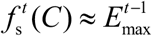.

### 2.2 Modal exploitation

In the modal exploitation phase, a distance-based scoring function is designed to accelerate the sampling of more reasonable conformations in the given basins. The distance-based scoring function is computed by:

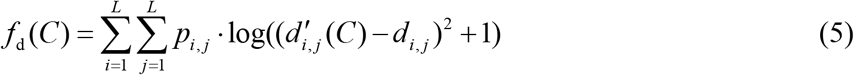

where *L* is the length of the protein sequence, 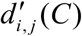 is the distance between *C_β_* atoms (*C_α_* atoms for glycine) in the *i*-th and *j*-th residues of the target conformation *C, d_i,j_* is the predicted distance of the residue pair (*i, j*) in the distance distribution, and *p_i,j_* is the probability that the predicted distance of the residue pair (*i, j*) is *d_i,j_*.

The seed conformations are optimized under the guidance of the original energy function and the distance-based scoring function to obtain the conformation closer to the native structure. *T* MMC trajectories are performed with *T* seed conformations as the initial, and the sampling range in each trajectory is limited to the basin region where the seed conformation is located. As illustrated in Figure 5, if the conformation escapes from Basin 2, the sampling move is rejected directly. Otherwise, the sampling move is accepted probabilistically (see flowchart in Supplementary Figure S3). The acceptance probability *P*_ace_(*C*) is defined as:

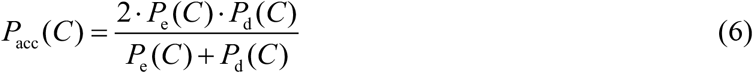

where *P*_e_ (*C*) and *P*_d_ (*C*) are the Boltzmann acceptance probabilities of the original energy function and the distance-based scoring function, respectively.

**Fig. 5.**
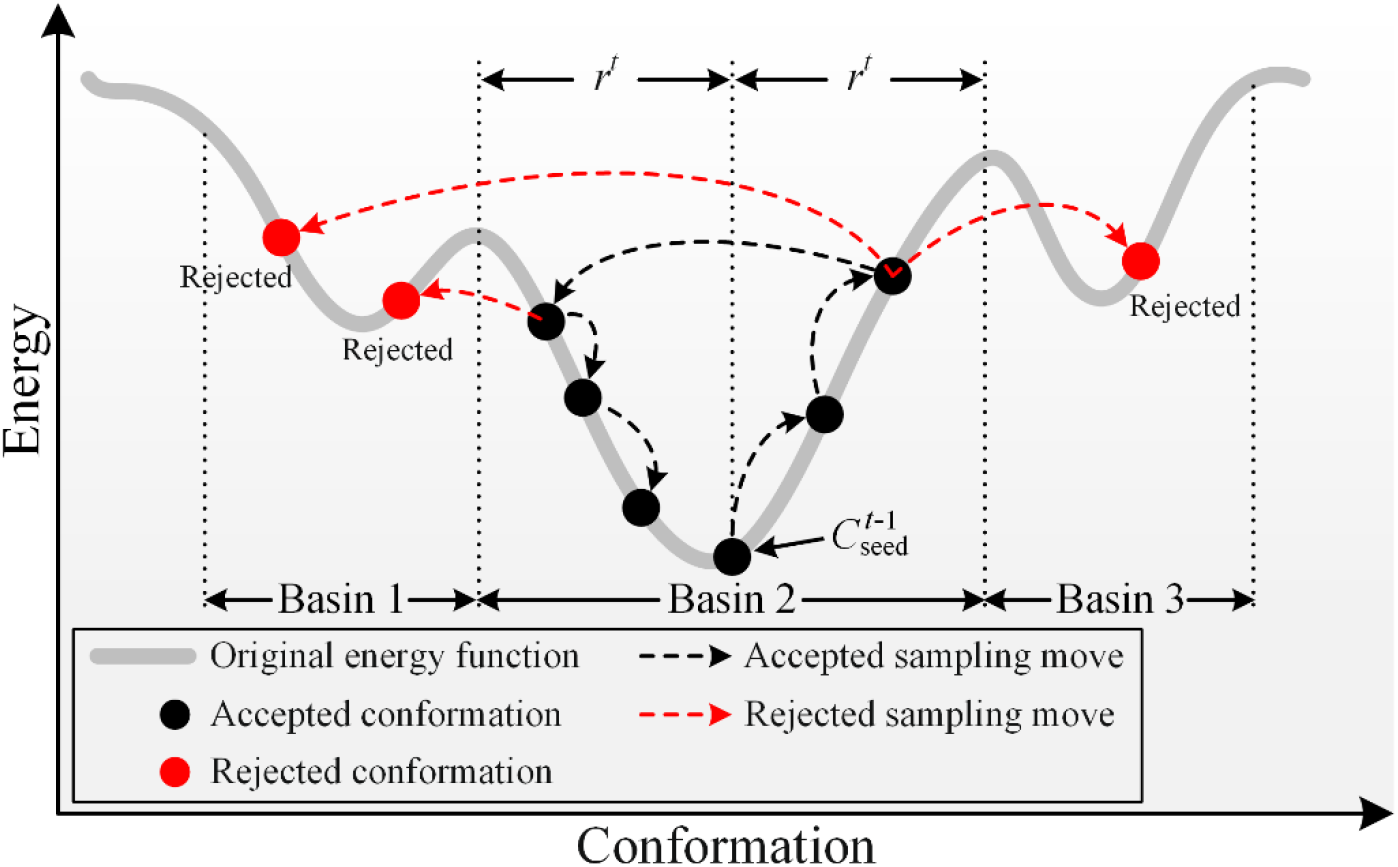
Schematic of modal exploitation. Starting from the seed conformation, MMC simulation is run again under the original energy function, supplemented by the distance-based scoring function. The sampling range is the basin where the seed conformation is located (Basin 2). In the sampling, the conformations (red dots) that jump out of the Basin 2 are rejected directly, and the red arrows indicate the rejected sampling moves. The conformations (black dots) staying in the Basin 2 are accepted with probability, and the black arrows are the accepted sampling moves.

## 3 Result and discussion

In CASP, all groups submit a maximum of five models per target, and are instructed that most emphasis in the assessments will be placed on the model they designate as “model 1” (intended to be the most accurate model) (Moult *et al.*, 2018). Therefore, we set the number of trajectories *T =* 5 in SNfold to generate five models for each target, and set the *increase*_ *cycles =* 10 in Rosetta ClassicAbinitio protocol. The other parameter settings are shown in Supplementary Table S1. The root-mean-square-deviation (RMSD) and template modeling score (TM-score) (Zhang and Skolnick, 2004; Xu and Zhang, 2010) are used to evaluate the quality of models.

### 3.1 Dataset

In this study, the performance of SNfold is tested over 300 benchmark proteins systematically selected from the PDB. The length of these proteins ranges from 52 to 199 residues, with < 30% sequence identity to each other (Supplementary Table S2). Firstly, 243,819 proteins with known structures from the SCOPe 2.07 (Fox *et al.*, 2014; Chandonia *et al.*, 2019) are clustered by CD-HIT (Li and Godzik, 2006; Huang *et al.*, 2010) with a 30% sequence identity cutoff, and result in 11,198 proteins. Then, 2,481 proteins are obtained after excluding the multidomain proteins and the proteins with a length of < 50 or >200 from the 11,198 proteins. Lastly, 300 proteins are selected from the 2,481 remaining proteins according to their length diversity as the benchmark set. To further test the performance of SNfold, we compare it with four state-of-the-art servers on 24 CASP13 FM targets. The length of the 24 FM targets varies from 41 to 354 residues (Supplementary Table S3).

### 3.2 Results of benchmark set

SNfold is compared with two well-known *ab initio* protein structure prediction methods, Rosetta and C-QUARK, on the benchmark set of 300 proteins. For the fairness of comparison, the distance-based scoring function (Equation (5)) is added to Rosetta, and the same conformation acceptance probability (Equation (6)) is adopted as that in SNfold. Rosetta restrained by distance is named as “Rosetta-dist”. Here, 500 independent trajectories are run using Rosetta’s ClassicAbinitio protocol, denoted as Rosetta-dist(500). In each trajectory, *increase cycles* is set to 1, considering the limits of our computational capacity available. The centroid models of the first five clusters clustered by SPICKER (Zhang and Skolnick, 2004) using all decoys are considered as the final models. The results of C-QUARK are predicted by its online server (https://zhanglab.ccmb.med.umich.edu/C-QUARK/), and the fragments which come from the protein with sequence identity to the target > 30% are removed.

The average results of the first models of SNfold, Rosetta-dist(500) and C-QUARK on the benchmark set are shown in Table 1, and the detailed results of each protein are shown in Supplementary Table S4. The average TM-score of SNfold’s first models is 0.597, which is 8.7% higher than that (0.549) of Rosetta-dist(500) and comparable with that (0.627) of C-QUARK. SNfold correctly folds (i.e., TM-score ? 0.5) 231 out of 300 targets, accounting for 77.0% of the total, an increase of 12.7% compared with Rosetta-dist(500). The head-to-head comparisons of SNfold with Rosetta-dist(500) and C-QUARK on the benchmark set are shown in Figure 6. SNfold achieves a higher TM-score than Rosetta-dist(500) and C-QUARK on 253 and 119 proteins, respectively. In addition, the number of the energy function evaluations (FE) is used to evaluate the computational cost. The FE of Rosetta-dist(500) is 1.51 × 10^7^, which is ~10 times higher than that (1.14 × 10^6^) of SNfold. The improved performance of SNfold is mainly due to the sampling-guided energy function, which reduces the re-sampling of the explored basins. As an example shown in Figure 7(a), SNfold can obtain native-like models with fewer MMC trajectories compared to Rosetta-dist(500). Figure 7(b) reports an example with inaccurate energy function that SNfold obtains the model with a higher accuracy than that of Rosetta-dist(500). Therefore, the basin where the native structure may be located is discovered because the previously explored basins are filled by the derating function.

**Fig. 6.**
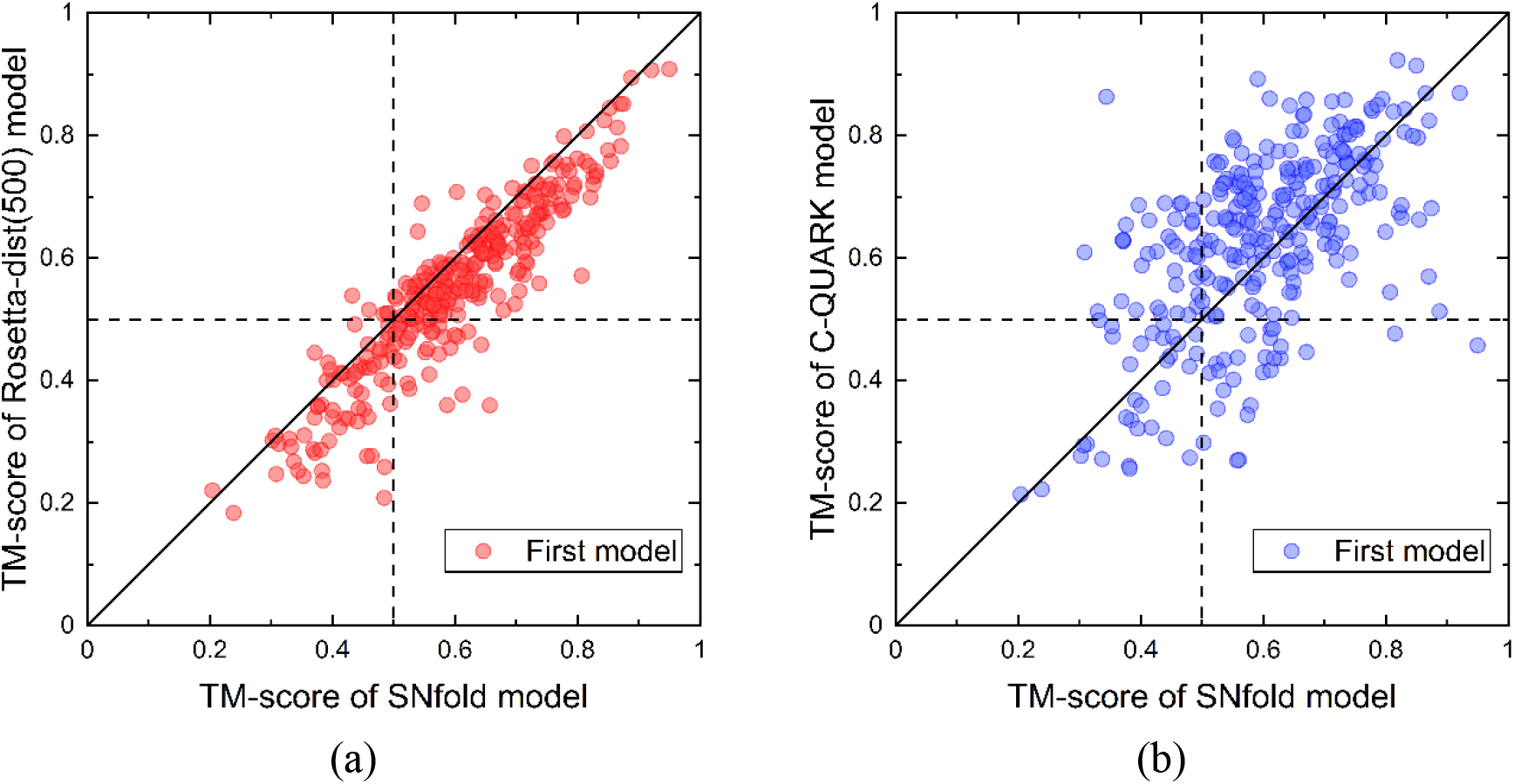
Summary of models generated by SNfold, Rosetta-dist(500) and C-QUARK. (a) Results of the head-to-head comparison between the first models predicted by SNfold and Rosetta-dist(500). (b) Results of the head-to-head comparison between the first models predicted by SNfold and C-QUARK.

**Fig. 7.**
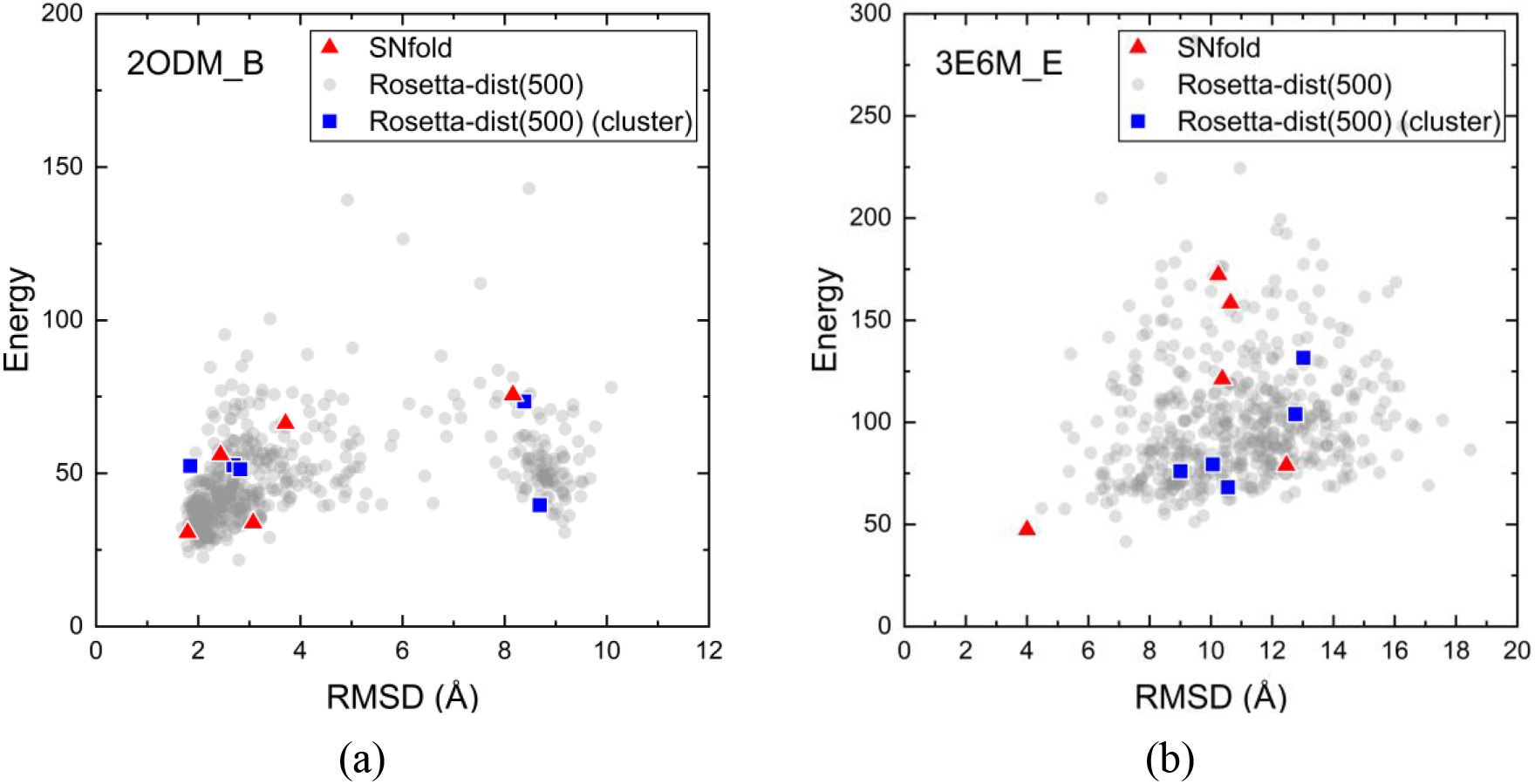
Two illustrative examples of the scatter plots of the models generated by SNfold and Rosetta-dist(500). (a) 2ODM_B. (b) 3E6M_E. The red dots represent the five models predicted by SNfold, the gray dots refer to the 500 models generated by 500 trajectories of Rosetta-dist(500), and the blue dots correspond to the five final models predicted by Rosetta-dist(500), which are clustered from the 500 models.

**Table 1:** Average results of SNfold, Rosetta-dist(500) and C-QUARK on the benchmark set. Correctly Fold represents the number of models with TM-score ≥ 0.5. FE is the number of the energy function evaluations, which is used to evaluate the computational cost.

| Method | RMSD ($\text{\AA}$) | TM-score | Correctly Fold | FE |
| --- | --- | --- | --- | --- |
| SNfold | 6.900 | 0.597 | 231/300 | $1.14 \times 10^6$ |
| Rosetta-dist(500) | 6.988 | 0.549 | 205/300 | $1.51 \times 10^7$ |
| C-QUARK | 6.701 | 0.627 | 242/300 | NA |

The average TM-score of the three methods for targets with different lengths is compared to verify the relationship between the accuracy of the prediction model and its length8(Figure 8). For targets with length of < 100, SNfold’s average TM-score is 0.622, which is 8.6% and 5.8% higher than those of Rosetta-dist(500) (0.573) and C-QUARK (0.588), respectively. SNfold (0.624) improves by 9.1% and 1.3% compared to Rosetta-dist(500) (0.572) and C-QUARK (0.616), respectively, when the target length < 120. For targets with length of < 150, SNfold’s TM-score improves by 8.1% over Rosetta-dist(500) (0.569) and is comparable with C-QUARK (0.615 vs. 0.630). For targets with length of > 150, SNfold’s TM-score is lower than that of C-QUARK because of several possible reasons. Firstly, C-QUARK contains an improved meta-method, NeBcon (He *et al.*, 2017), which combines multiple state-of-the-art contact predictors, including several deeplearning-based methods, such as ResPRE (Li *et al.*, 2019a,b). Secondly, C-QUARK involves REMC simulation whose computational cost may be much greater than that of SNfold, which results in more potential conformations can be sampled. Thirdly, SNfold is based on Rosetta which can generate high-resolution models for some small proteins, especially those < 100 residues in length (Bradley *et al.*, 2005). However, the accuracy is relative lower for proteins with > 150 residues as the conformational space is vastly increased.

**Fig. 8.**
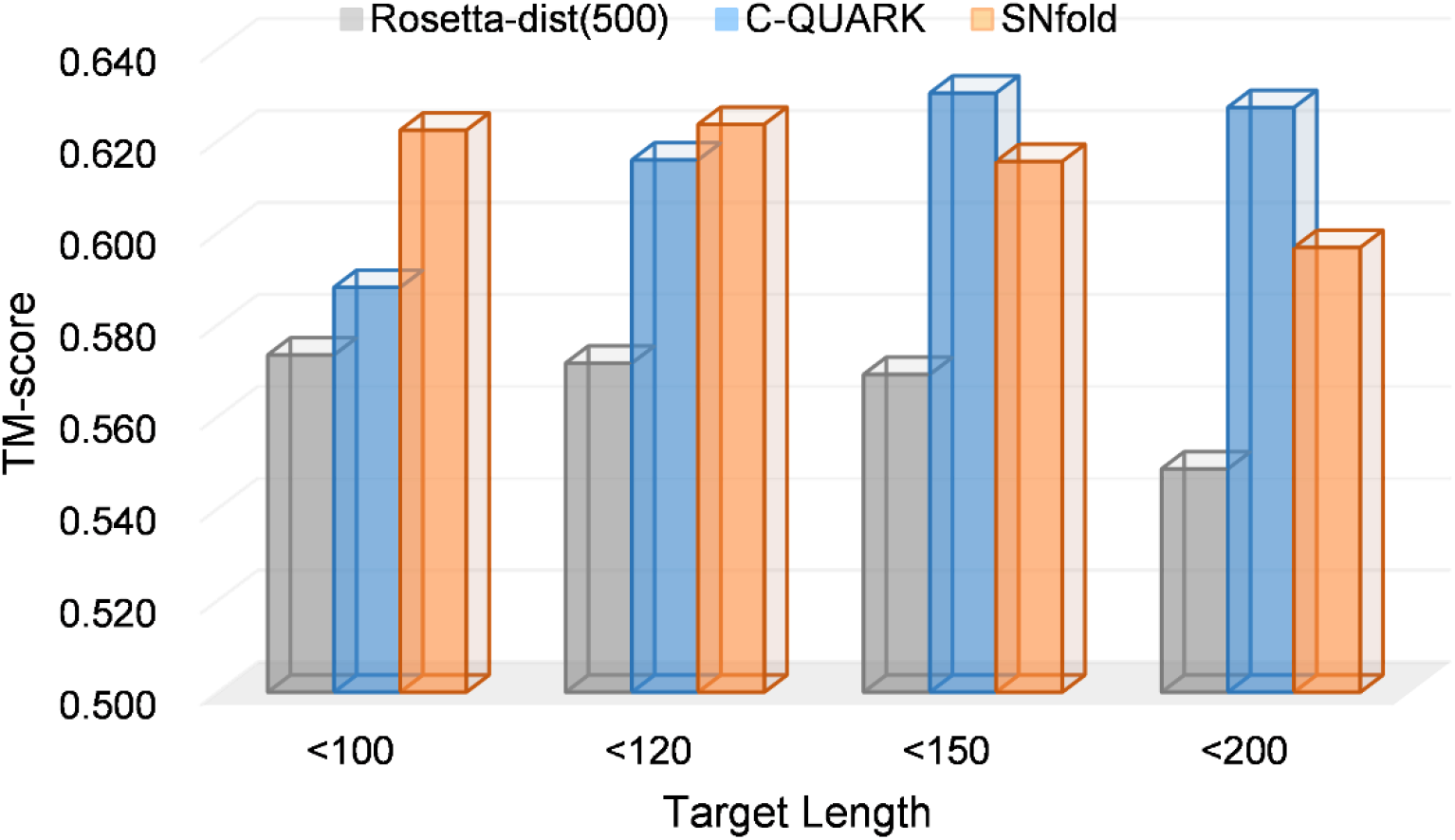
Comparison of SNfold, Rosetta-dist(500) and C-QUARK in protein groups with different length ranges.

### 3.3 Analysis of conformation sampling efficiency

To verify whether SNfold can reduce re-sampling of previously explored basins, we only run the modal exploration phase of SNfold, named as SNfold-exploration. For comparison, the same number of MMC trajectories are independently run by Rosetta. The two methods are compared over the 300 test proteins using the same computational cost, and neither uses the distance constraint. Here, a metric called retry rate (*η*_retry_) is defined to measure the extent to which the conformation repeatedly enters the explored basins during the sampling process:

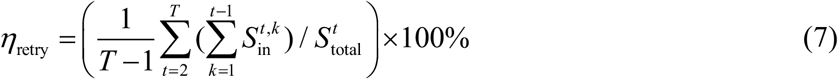

where *T* = 5 is the number of trajectories; 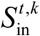 is the number of the accepted conformations of the *t*-th trajectory that re-enter the basin explored by the *k*-th trajectory; and 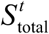 is the number of all accepted conformations in the *t*-th trajectory. The detailed description is shown in Supplementary Figure S4. *η*_retry_ reflects the efficiency of conformation sampling. The smaller *η*_retry_, the lesser the re-sampling of the previously explored basins and the higher the efficiency of conformation sampling.

The results of SNfold-exploration and Rosetta are shown in Table 2 and Supplementary Figure S5. *η*_retry_ of SNfold-exploration is 3.36%, which is significantly lower than that (65.74%) of Rosetta (*P*-value = 6.08 × 10^-51^). This finding reflects that randomly starting multiple independent MMC trajectories is indeed inefficient, because there is a high probability of re-entering the previously explored basins in the subsequent sampling. Compared with Rosetta, SNfold-exploration achieves a lower retry rate because the previously explored basins are filled by the derating function. Therefore, SNfold can reduce the re-sampling of previously explored basins and improve the efficiency of conformation sampling.

**Table 2.** Average retry rate (*η*_retry_) of SNfold-exploration and Rosetta on the benchmark set. The last two columns are the results of the Wilcoxon signed-rank test calculated with *η*_retry_.

| Method | $\eta_{\text{retry}}$ (%) | $P$-value | Significance |
| --- | --- | --- | --- |
| SNfold-exploration | 3.36 | NA | NA |
| Rosetta | 65.74 | $6.08 \times 10^{-51}$ | + |

Figure 9 shows the conformational distribution of SNfold-exploration and Rosetta for the target 1ELW_A. There are two “funnels” with densely distributed conformations (the red circles in Figure 9(a)(b)). This indicates that at least two low-energy basins likely exist on the energy surface. The five models produced by Rosetta are all trapped in “funnel 2”, which may be caused by the repeated exploration of the low-energy region (Figure 9(a)). For SNfold-exploration (Figure 9(b)), in addition to the four conformations in “funnel 2”, another conformation (model 2) exists in “funnel 1” as it uses the derating function. The RMSD and TM-score of SNfold-exploration’s model 2 are 1.50 Å and 0.88, respectively, which are better than those of all five models generated by Rosetta ((11.93 Å, 0.45), (8.48 Å, 0.55), (10.82 Å, 0.52), (10.77 Å, 0.47) and (10.93 Å, 0.44)). The model 1 generated by SNfold-exploration is located in “funnel 2” and the derating function fills the basin where model 1 is located. Therefore, in the second trajectory, the re-sampling of the basin is reduced (the blue asterisks in “funnel 2” are less), and sampling of other basins is increased (the blue asterisks in “funnel 1” are more). This result also indicates that SNfold can increase the possibility of exploring other basins where the native structure may be located, thereby improving the reliability of prediction models.

**Fig. 9.**
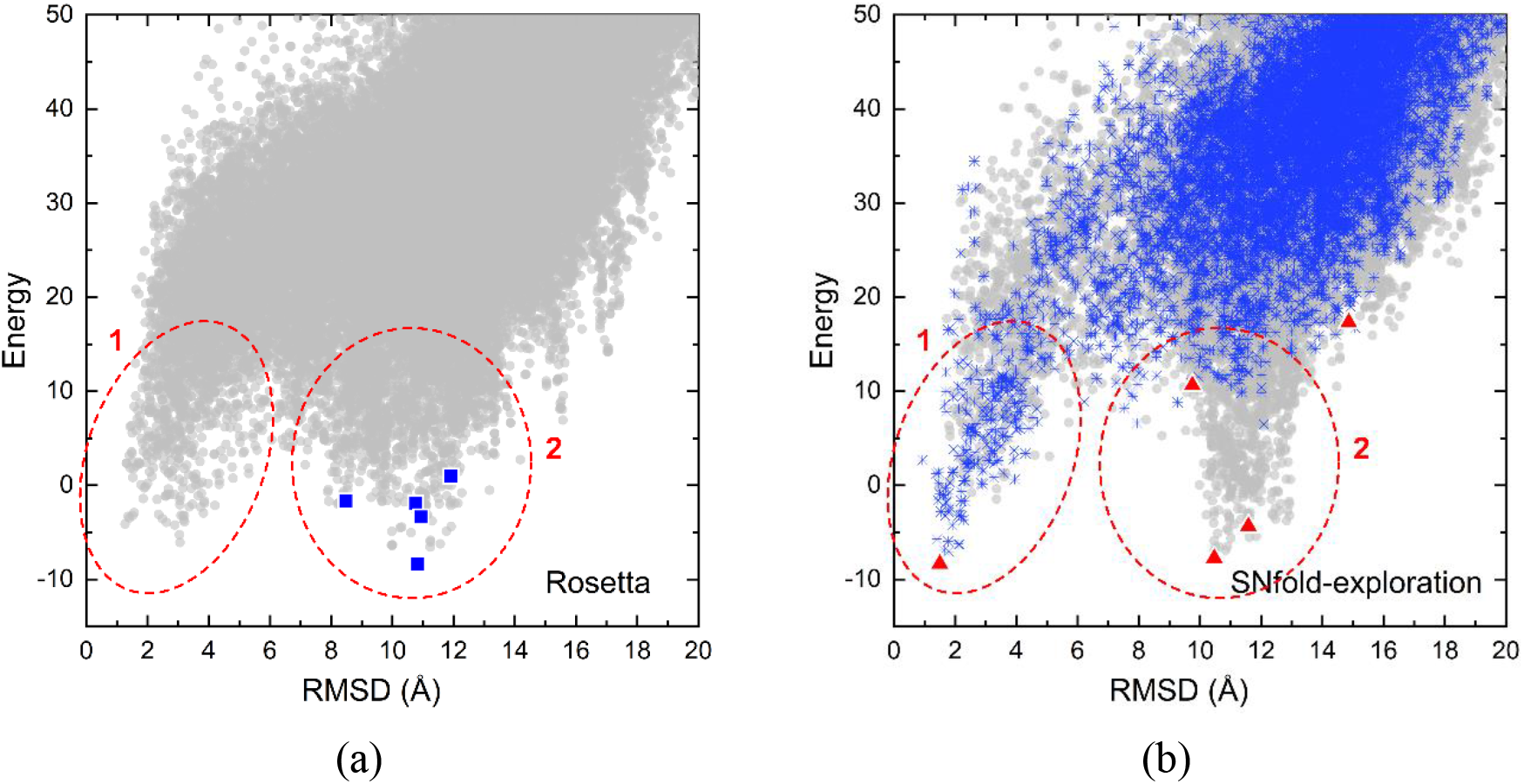
Conformational distribution of SNfold-exploration and Rosetta for 1ELW_A. (a) Conformational distribution of Rosetta. The gray dots represent all the accepted conformations sampled in five trajectories of Rosetta, and the blue square denotes the lowest energy conformation in each trajectory. (b) Conformational distribution of SNfold-exploration. The gray dots indicate all the accepted conformations sampled in five trajectories of SNfold- exploration, the blue asterisks refer to the accepted conformations sampled in the second trajectory of SNfold- exploration, and the red triangle corresponds to the lowest energy conformation in each trajectory.

In order to further compare the computational cost of SNfold and Rosetta-dist, here we increase the trajectories of Rosetta-dist to 1,000, annotated as Rosetta-dist(1000). The comparison between SNfold and Rosetta-dist(1000) is presented in Table 3 and Figure 10, and the detailed results are listed in Supplementary Table S5. In Table 3, the average RMSD of SNfold and Rosetta-dist(1000) are 5.984 Å and 5.593 Å, respectively. The average TM-score of SNfold is 0.614, which is comparable with that of Rosetta-dist(1000) (*P*-value = 0.265). However, the FE of SNfold is less than that of Rosetta-dist(1000), which is about 1/200 of the latter. This shows that the computational efficiency of SNfold on the given test set is more than 100 times higher than that of Rosetta-dist(1000) with almost no loss of accuracy.

**Fig. 10.**
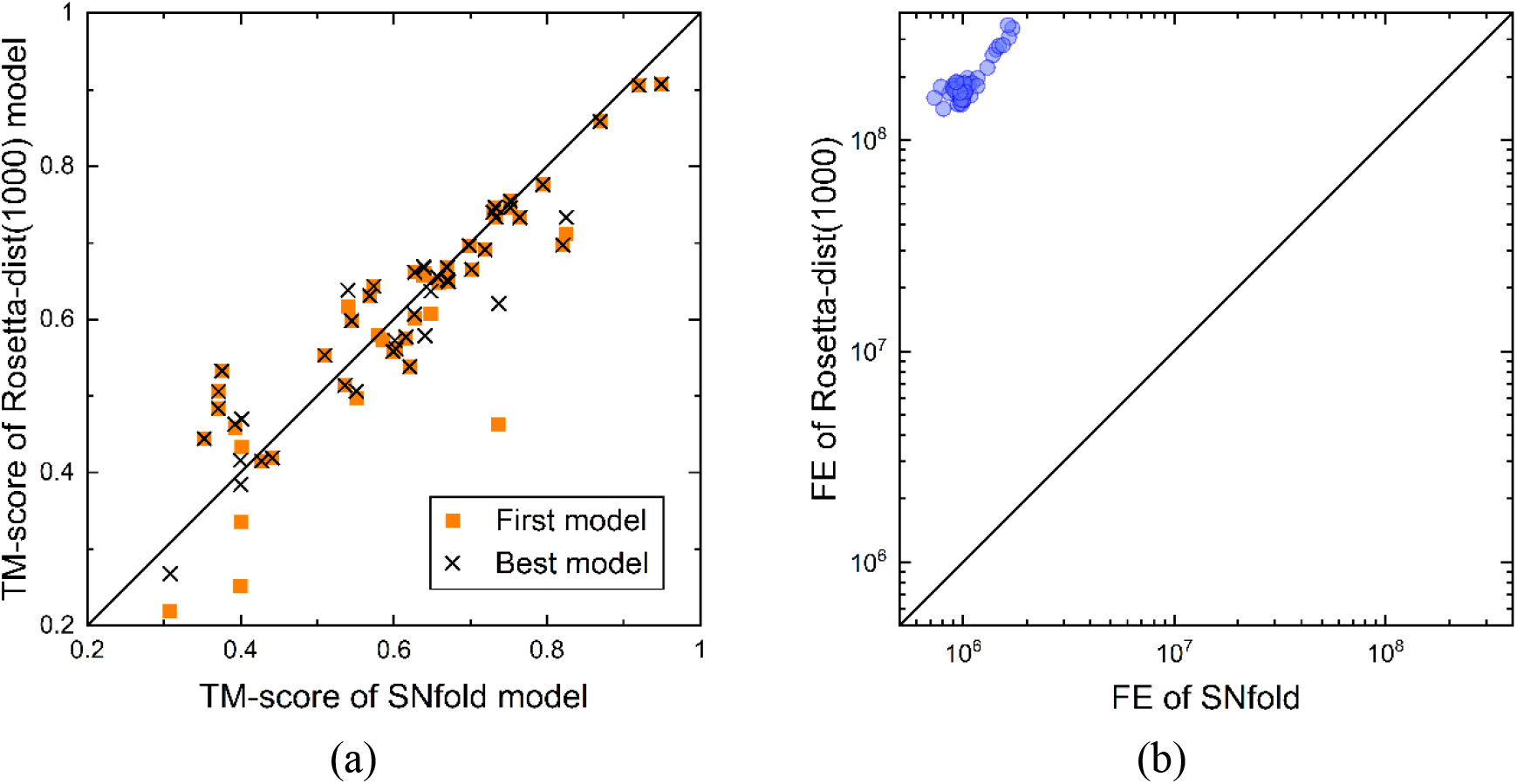
Comparison of SNfold and Rosetta-dist(1000). (a) Head-to-head comparison between the TM-score of the first models and the best models by SNfold and Rosetta-dist(1000). (b) Comparison between FE of SNfold and Rosetta-dist(1000).

**Table 3.** Average prediction accuracy and number of the energy function evaluations (FE) of SNfold and Rosetta- dist(1000) on 50 targets randomly selected from the benchmark set. *P*-value is calculated with the TM-score of SNfold and Rosetta-dist(1000).

| Method | RMSD (Å) | TM-score | *P*-value | FE |
| --- | --- | --- | --- | --- |
| SNfold | 5.984 | 0.614 | NA | $1.06 \times 10^6$ |
| Rosetta-dist(1000) | 5.593 | 0.605 | 0.265 | $1.91 \times 10^8$ |

### 3.4 Component analysis

In order to verify the effect of modal exploration, SNfold is compared with SNfold-exploitation (SNfold without modal exploration) on the benchmark set. The results are summarized in Table 4, and the detailed results are presented in Supplementary Table S4. The comparison of the performance of SNfold and SNfold-exploitation is illustrated in Figure 11. The average RMSD and TM-score of the first model generated by SNfold-exploitation are 8.676 Å and 0.501, respectively. That is, the average RMSD of SNfold is decreased by 20.5%, and the average TM-score is increased by 19.2% when the exploration process is added. The RMSD of SNfold is lower than that of SNfold- exploitation on 235 proteins, and the TM-score of the former is higher than that of the latter on 278 proteins. The number of models correctly folded by SNfold is 54.0% more than that by SNfold- exploitation. The results indicate the effectiveness of modal exploration, which can further improve the accuracy of the prediction models compared with that of using the distance-based scoring function alone.

**Fig. 11.**
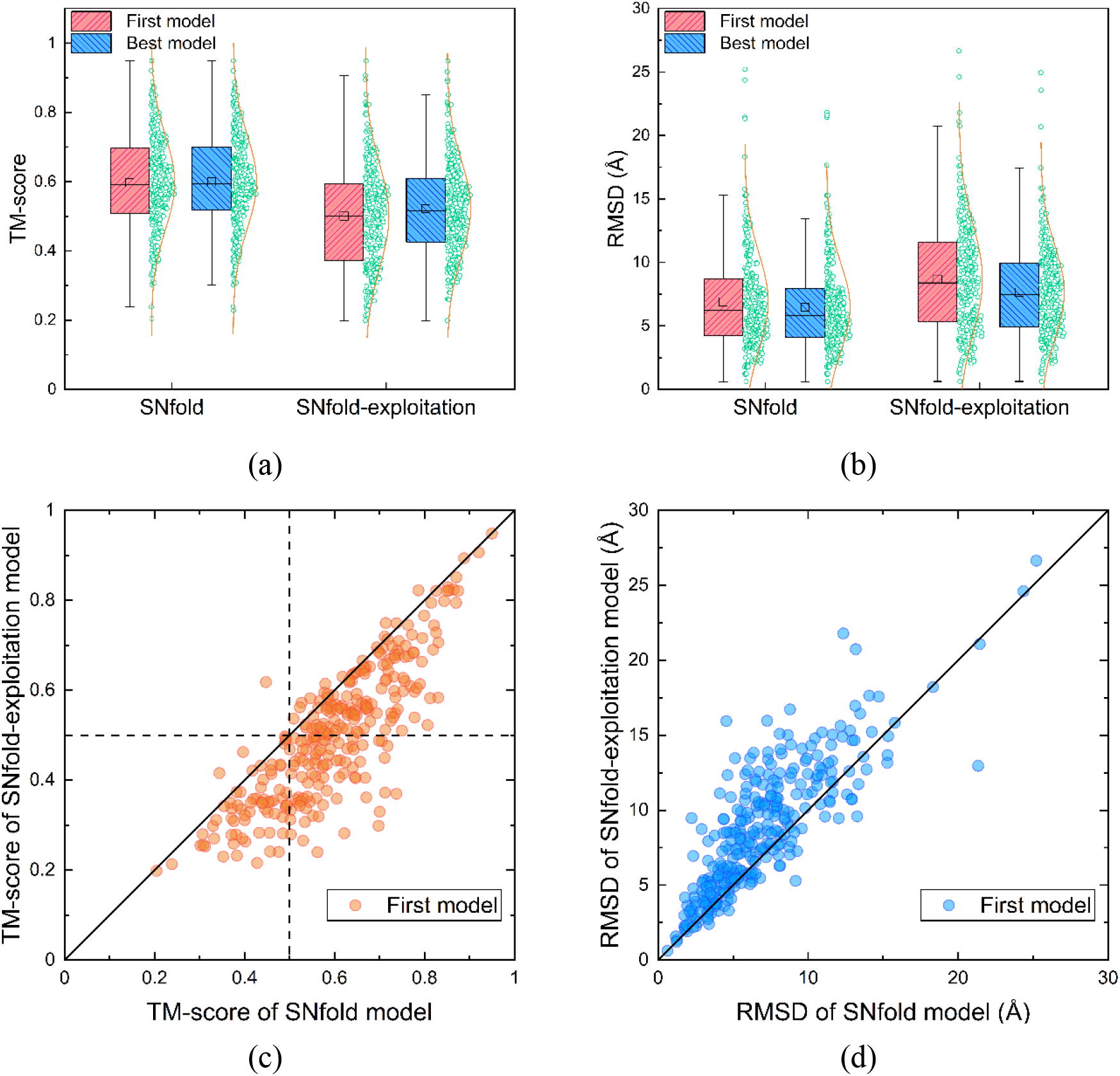
Comparison of the prediction performance between SNfold and SNfold-exploitation on the benchmark set. (a) Boxplot and distribution for TM-score of the first models and the best models by SNfold and SNfold-exploitation, respectively. (b) Boxplot and distribution for RMSD of the first models and the best models by SNfold and SNfold- exploitation, respectively. (c) Head-to-head comparison between TM-score of the first models by SNfold and SNfold-exploitation. (d) Comparison between RMSD of the first models by SNfold and SNfold-exploitation.

**Table 4.** Average prediction results of SNfold and SNfold-exploitation on the benchmark set.

| Method | RMSD (Å) | TM-score | Correctly Fold |
| --- | --- | --- | --- |
| SNfold | 6.900 | 0.597 | 231/300 |
| SNfold-exploitation | 8.676 | 0.501 | 150/300 |

### 3.5 Results of CASP13 targets

We also tested our method on 24 FM targets from CASP13, and compared it with four state-of-the-art methods of server group in CASP13, i.e., QUARK (Zheng *et al.*, 2019), BAKER-ROSETTASERVER, RaptorX-DeepModeller (Xu and Wang, 2019) and MULTICOM_CLUSTER (Hou *et al.*, 2019) (Table 5 and Supplementary Table S6). The results of QUARK, BAKER-ROSETTASERVER, RaptorX-DeepModeller and MULTICOM_CLUSTER are from the CASP official website (https://predictioncenter.org/casp13/results.cgi). On the 24 CASP13 FM targets, the average TM-score of the first model by SNfold is 0.461, which is 10.3% and 24.3% higher than that of BAKER-ROSETTASERVER (0.418) and MULTICOM_CLUSTER (0.371), respectively. It is comparable with QUARK (0.496) and RaptorX-DeepModeller (0.491). This is probably because SNfold does not use templates information and only runs five MMC trajectories.

**Table 5.** Average prediction results of SNfold, QUARK, BAKER-ROSETTASERVER, RaptorX-DeepModeller and MULTICOM_CLUSTER on 24 FM targets from CASP13.

| Group (Method) | TM-score | Correctly Fold |
| --- | --- | --- |
| SNfold | 0.461 | 8/24 |
| QUARK | 0.496 | 12/24 |
| BAKER-ROSETTASERVER | 0.418 | 6/24 |
| RaptorX-DeepModeller | 0.491 | 13/24 |
| MULTICOM_CLUSTER | 0.371 | 5/24 |

The TM-score of the first model for each target by the five methods is illustrated in Figure 12. SNfold correctly folds 8 models with TM-score ≥ 0.5 amongst 24 FM targets and obtains the highest TM-score on 8 targets. An example of the improved performance of SNfold is shown in Figure 13(a) for target T0957s2-D1; the TM-score of this model is 0.670, whereas the values obtained by other methods are: QUARK (0.53), BAKER-ROSETTASERVER (0.48), RaptorX-DeepModeller (0.65) and MULTICOM_CLUSTER (0.51). For target T1008-D1 (Figure 13(b)), the first model of SNfold also obtains the highest TM-score (0.666), and is 90.3%, 18.9%, 137.9% and 75.3% higher than that of QUARK (0.35), BAKER-ROSETTASERVER (0.56), RaptorX-DeepModeller (0.28) and MULTICOM_CLUSTER (0.38), respectively. There are three targets with lengths > 300, i.e., T0950-D1, T0969-D1 and T1005-D1 which have 342, 354 and 326 residues, respectively. The accuracy of SNfold on the targets T0969-D1 and T1005-D1 is relatively poor compared with other methods. This may be due to the decreasing performance of SNfold with the increase in protein length or the inaccurate distance distribution predicted. However, there is an exception to the target T0950- D1 (Figure 13(c)). The first model predicted by SNfold has TM-score = 0.509, except RaptorX-DeepModeller (0.56), which is higher than other three methods: QUARK (0.44), BAKER- ROSETTASERVER (0.46) and MULTICOM_CLUSTER (0.22).

**Fig. 12.**
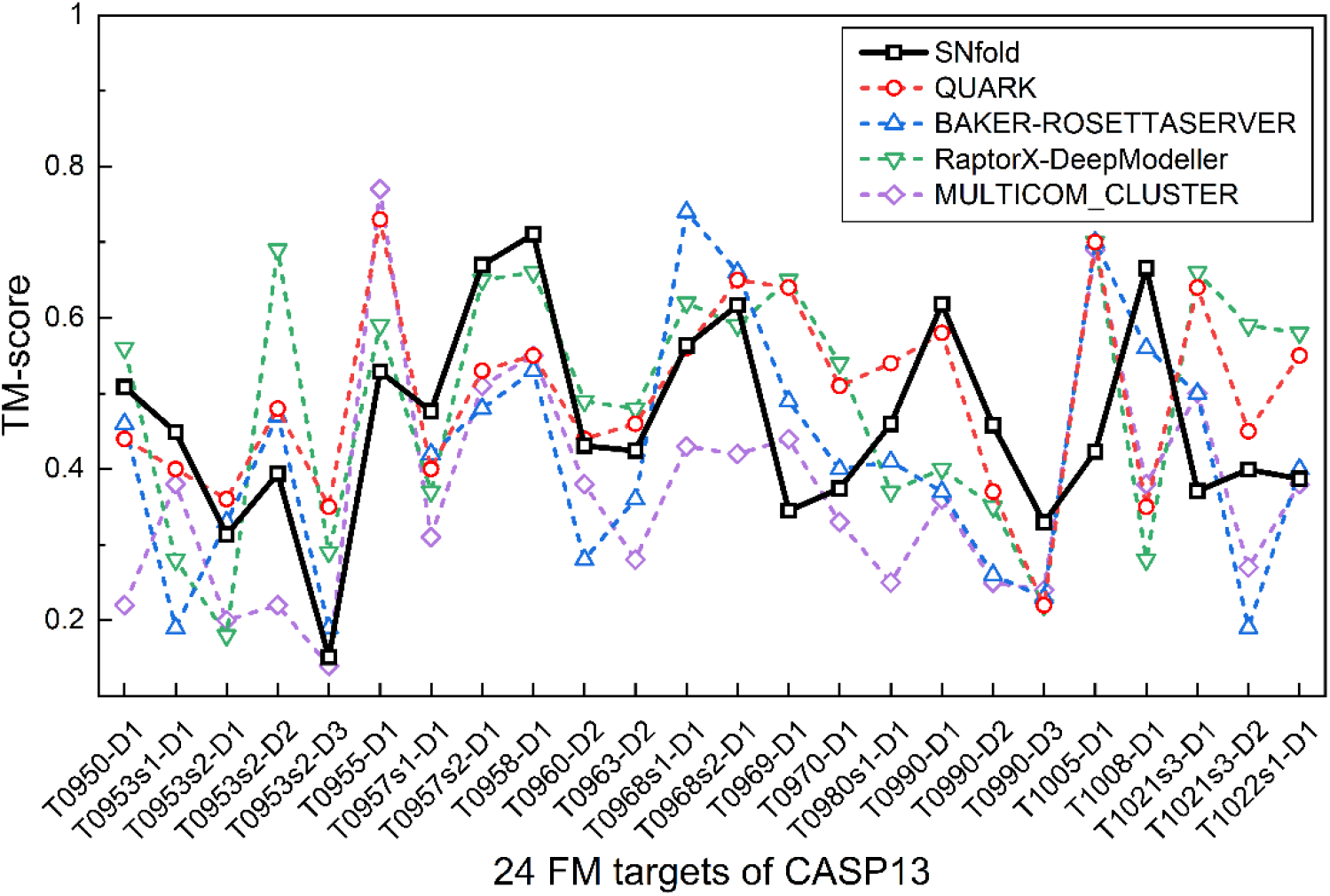
TM-score of the first models predicted by SNfold, QUARK, BAKER-ROSETTASERVER, RaptorX-DeepModeller and MULTICOM_CLUSTER for 24 FM targets from CASP13.

**Fig. 13.**
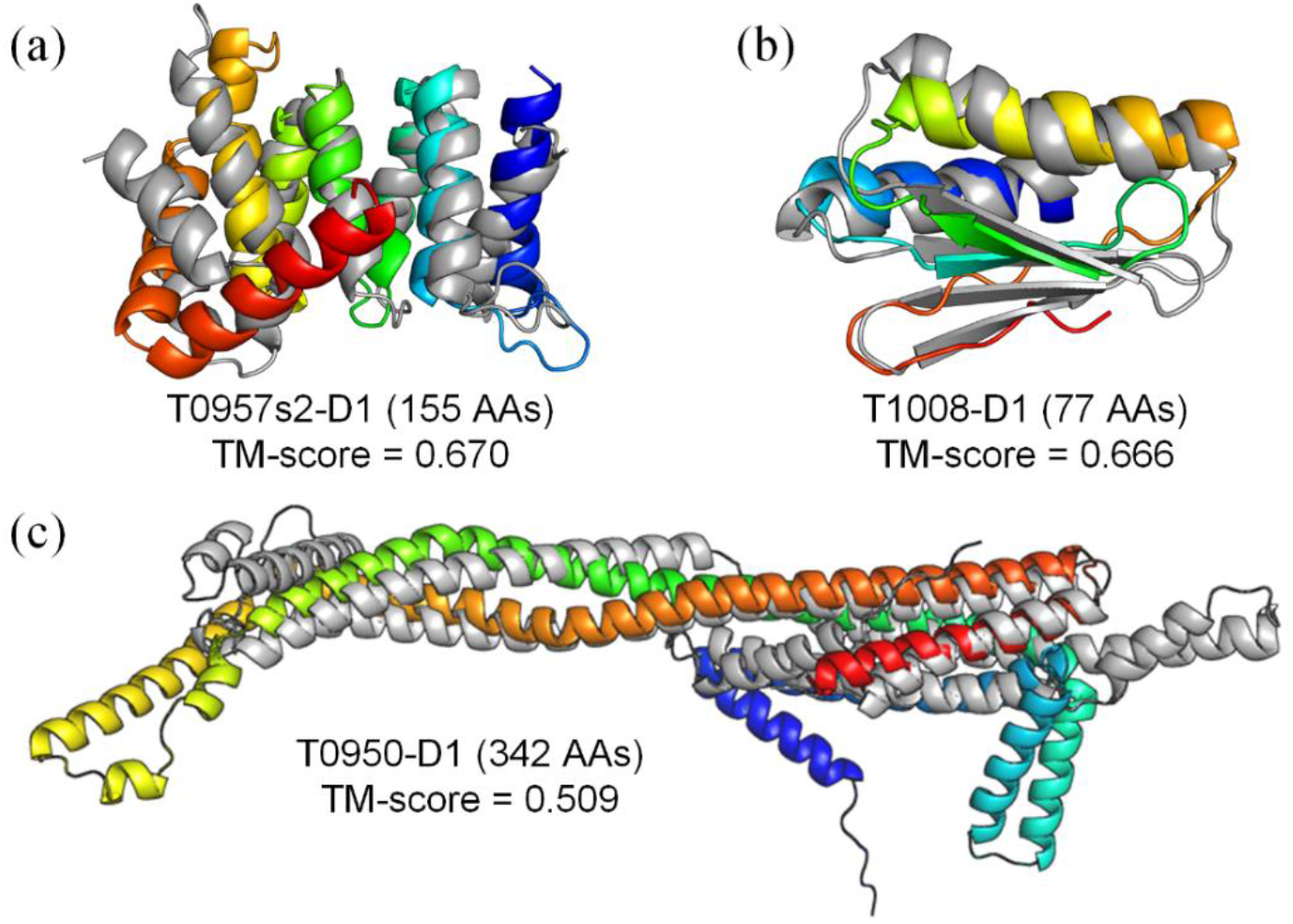
Superimposition between the first model (rainbow) by SNfold and the native structure (gray) for three CASP13 FM targets (T0957s2-D1, T1008-D1 and T0950-D1).

## 4 Conclusion

We have developed a sequential niche multimodal conformation sampling algorithm, SNfold, to improve the conformation sampling efficiency in protein structure prediction without loss of accuracy. In SNfold, a series of sampling-guided energy functions are constructed by the derating function designed from the previous sampling. With the sampling-guided energy functions, the sampling algorithm avoids the re-sampling of the explored basins and increases the likelihood of navigating potential basins where the native structure may be located. Meanwhile, a distance-based scoring function is designed to accelerate sampling with more reasonable structures in the previously explored basins.

SNfold is tested on 300 benchmark proteins and 24 FM targets from CASP13. Experimental results show that the accuracy of SNfold is comparable with Rosetta-dist and C-QUARK. SNfold correctly folds 231 out of 300 benchmark proteins. However, in terms of sampling efficiency, SNfold increases the computational efficiency by more than 100 times compared with Rosetta-dist on the test set. On the 24 CASP13 FM targets, SNfold is also comparable with the four top-ranked methods in the CASP13 server group.

As a plug-in conformation sampling algorithm, SNfold can be applied to other protein structure prediction methods. However, the performance of SNfold still remains to be improved for larger proteins due to the limitation of fragment assembly. Considering that fragment assembly contains native structure information, and geometric optimization has a stronger sampling ability for local basins, one way to alleviate this issue may be the combination of the fragment assembly and distance-based geometric optimization. In addition, trying to reveal the folding mechanism of proteins based on the idea in SNfold is also a direction of our follow-up exploration.

