## Supplementary data for "A Sequential Niche Multimodal Conformation Sampling Algorithm for Protein Structure Prediction"

### Supplementary Information

Yu-Hao Xia, Chun-Xiang Peng, Xiao-Gen Zhou and Gui-Jun Zhang

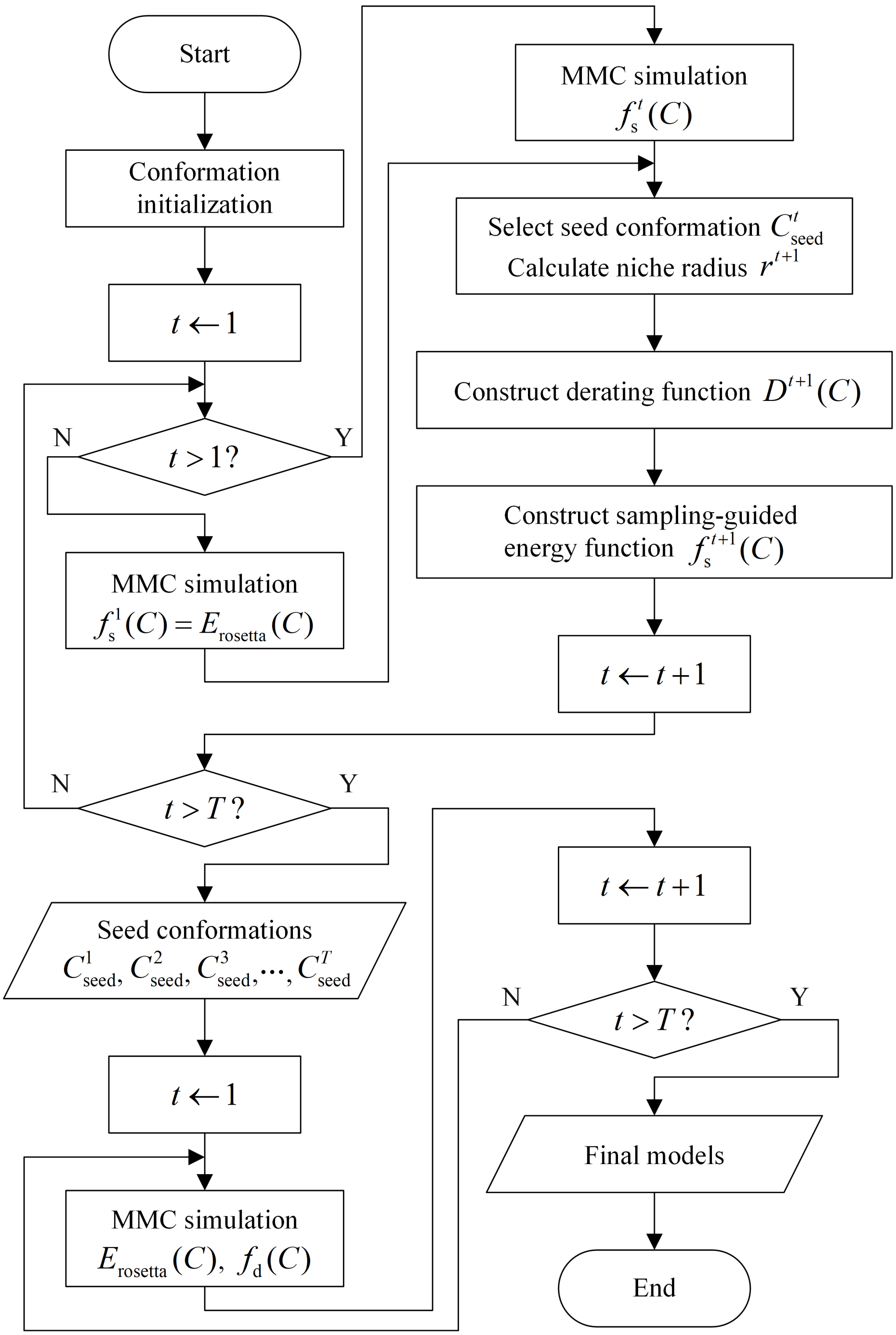


Figure S1: Flowchart of SNfold. $T$ is the number of trajectories; $C$ is the target conformation; $f_s^1(C)$ is the original energy function $E_{rosetta}(C)$; $f_s^t(C)$ is the sampling-guided energy function of the $t$-th trajectory; $r^{t+1}$ is the niche radius calculated by seed conformation $C_{seed}^t$; $D^{t+1}(C)$ is the derating function; and $f_d(C)$ is the distance-based scoring function.

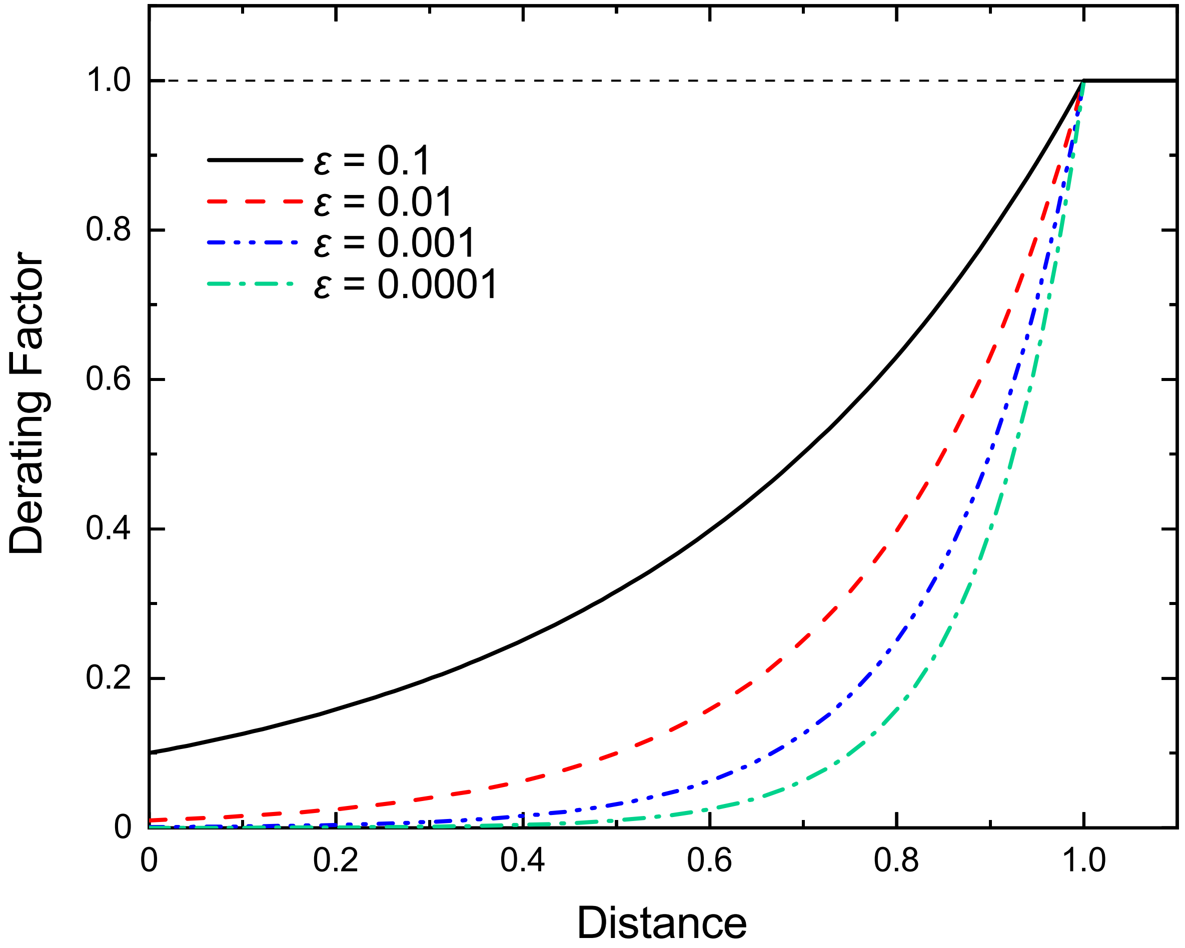


Figure S2: Derating curves for different values of $\epsilon$. The $x$-axis is the distance between the target conformation and the seed conformation; the $y$-axis is the derating factor. Here, the niche radius is assumed to be 1.

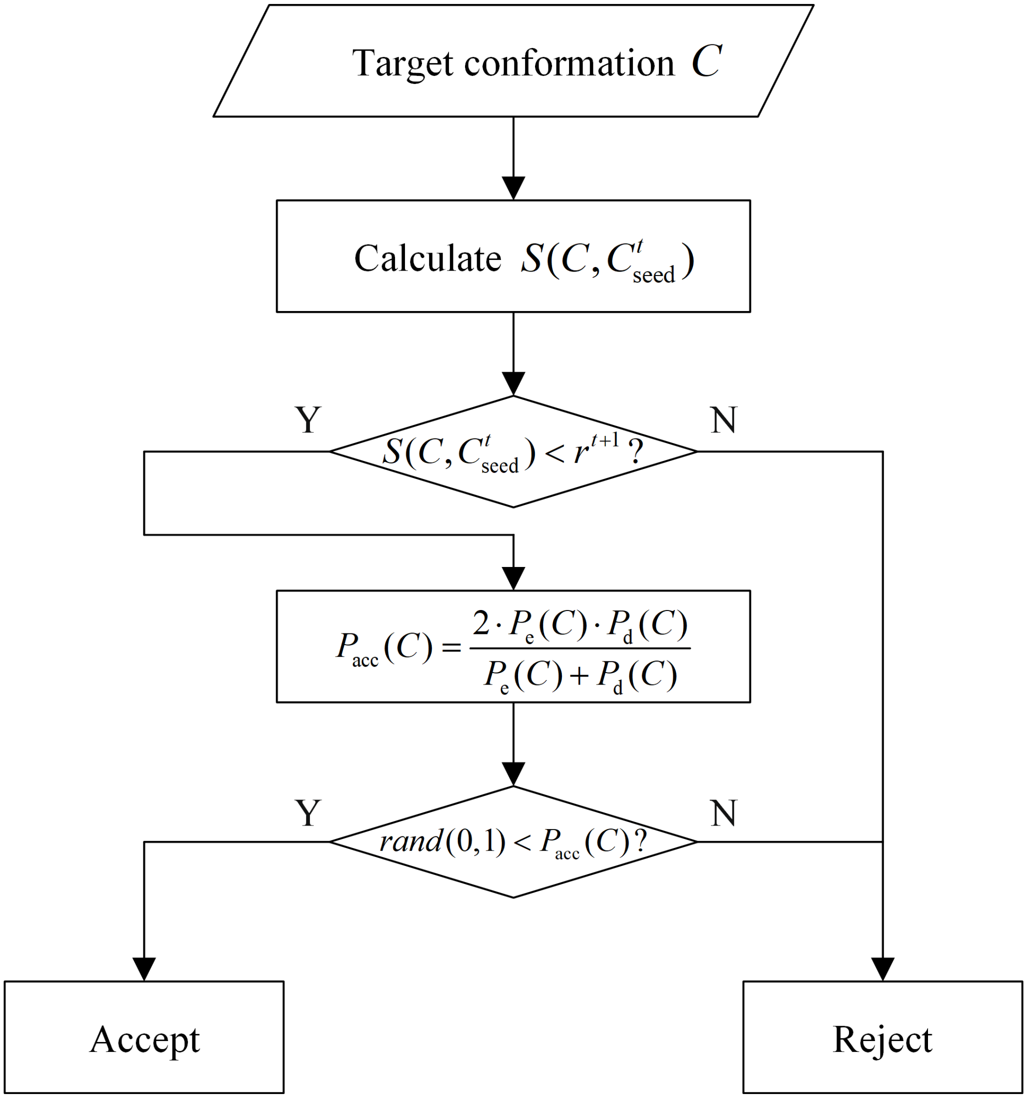


Figure S3: Flowchart of conformation selection strategy in the modal exploitation phase. $S(C, C^t_{\text{seed}})$ is the distance between the target conformation $C$ and the seed conformation $C^t_{\text{seed}}$; $r^{t+1}$ is the niche radius; $P_{\text{acc}}(C)$ is the acceptance probability for the target conformation $C$, where $P_{\text{e}}(C) = \min(\exp(-(E_{\text{rosetta}}(C) - E_{\text{rosetta}}(C'))/\beta_1), 1)$, $P_{\text{d}}(C) = \min(\exp(-(f_{\text{d}}(C) - f_{\text{d}}(C'))/\beta_2), 1)$, $C'$ is the last accepted conformation, $\beta_1$ and $\beta_2$ are the temperature scaling factors of the Boltzmann acceptance probability of the original energy function and the distance-based scoring function, respectively; $rand(0, 1)$ is a random number between 0 and 1.

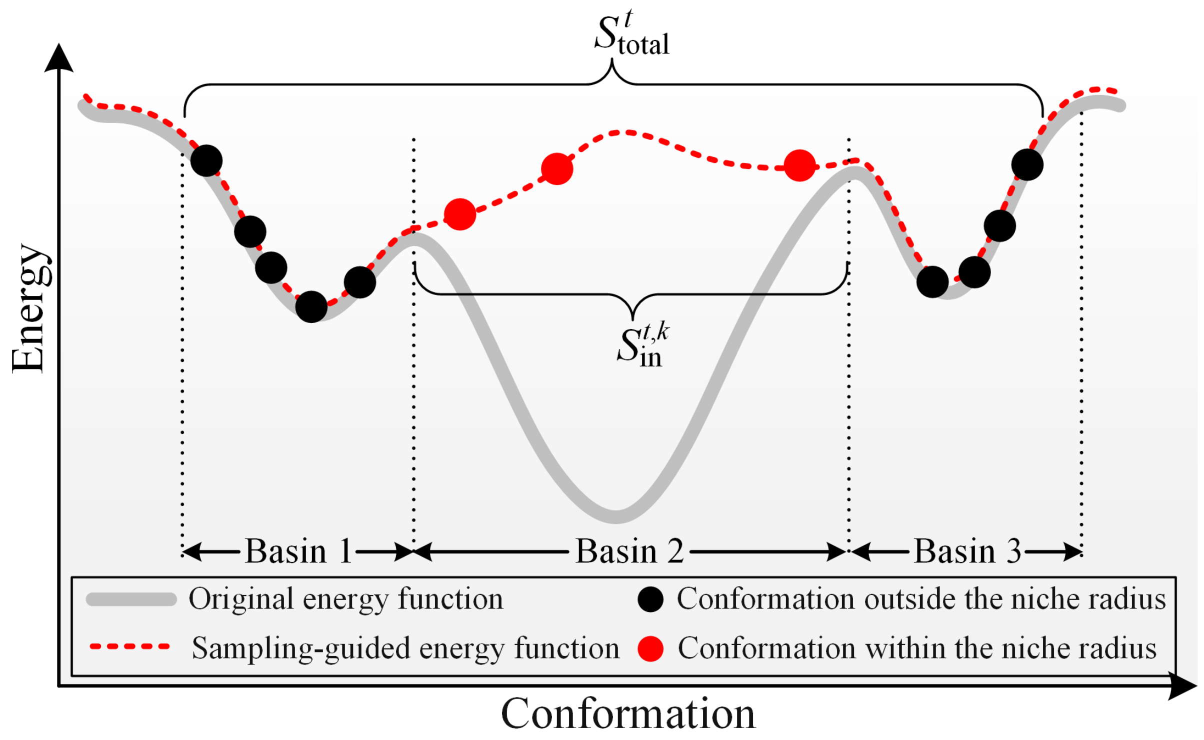


Figure S4: Schematic of retry rate ($\eta_{\text{retry}}$) definition. $S_{\text{in}}^{t,k}$ is the number of the accepted conformations of the $t$-th trajectory that re-enter the basin explored by the $k$-th trajectory (Basin 2); and $S_{\text{total}}^{t}$ is the number of all accepted conformations in the $t$-th trajectory.

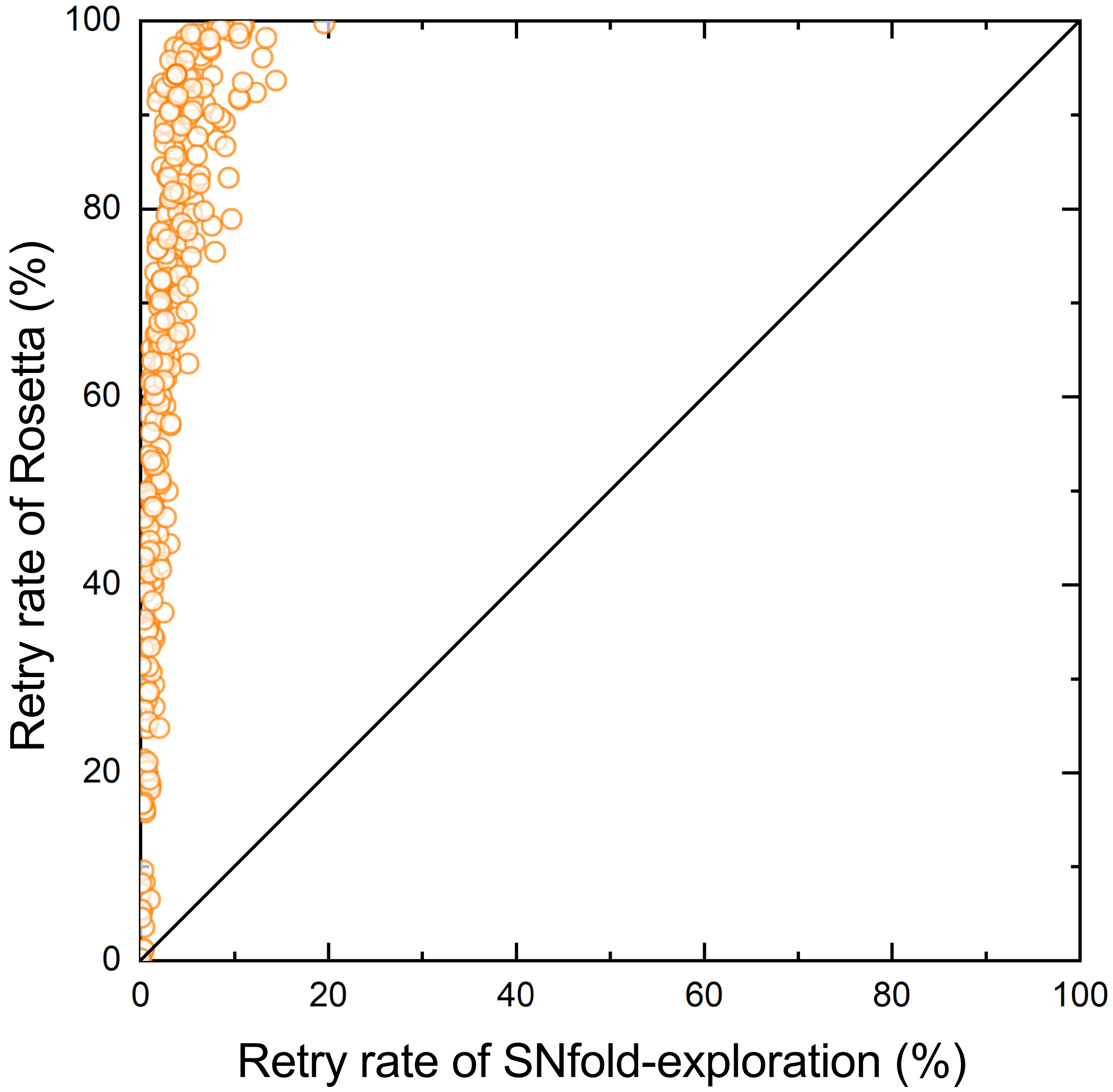


Figure S5: Results of the head-to-head comparison between the retry rate of SNfold-exploration and Rosetta.

Table S1: Parameter settings of SNfold. $T$ is the number of trajectories, which is also equal to the number of final prediction models; $\beta_1$ and $\beta_2$ are the temperature scaling factors of the Boltzmann acceptance probability of energy function and distance-based scoring function, respectively.

| Parameters of SNfold | | |
|---|---|---|
| Number of trajectories | $T$ | 5 |
| Derating minimum value | $\epsilon$ | 0.001 |
| Temperature scaling factor of $P_{\mathrm{e}}(C)$ | $\beta_1$ | 2 |
| Temperature scaling factor of $P_{\mathrm{d}}(C)$ | $\beta_2$ | 0.5 |

Table S2: Detailed information of 300 benchmark proteins used in the experiments of this study.

| Protein | Type | Size | Protein | Type | Size | Protein | Type | Size | Protein | Type | Size |
|---|---|---|---|---|---|---|---|---|---|---|---|
| 1A3A_C | $\alpha/\beta$ | 146 | 1A6L_A | $\alpha/\beta$ | 106 | 1A7D_A | $\alpha$ | 118 | 1A91_A | $\alpha$ | 79 |
| 1ABV_A | $\alpha$ | 105 | 1AHK_A | $\beta$ | 129 | 1AK6_A | $\alpha/\beta$ | 174 | 1AKP_A | $\beta$ | 114 |
| 1AP7_A | $\alpha$ | 168 | 1AUU_A | $\beta$ | 55 | 1AX8_A | $\alpha/\beta$ | 130 | 1B4R_A | $\beta$ | 80 |
| 1B4U_A | $\alpha$ | 132 | 1BE3_J | $\alpha$ | 62 | 1BGF_A | $\alpha/\beta$ | 124 | 1BGY_J | $\alpha$ | 62 |
| 1BJX_A | $\alpha/\beta$ | 110 | 1BUO_A | $\alpha/\beta$ | 121 | 1C03_A | $\alpha/\beta$ | 163 | 1C41_A | $\alpha/\beta$ | 165 |
| 1C9F_A | $\alpha/\beta$ | 87 | 1CDB_A | $\beta$ | 105 | 1CF7_B | $\alpha/\beta$ | 82 | 1CTO_A | $\beta$ | 109 |
| 1CXZ_B | $\alpha$ | 86 | 1D6T_A | $\alpha/\beta$ | 117 | 1D8B_A | $\alpha$ | 81 | 1DBF_A | $\alpha/\beta$ | 127 |
| 1DCF_A | $\alpha/\beta$ | 133 | 1DL6_A | $\beta$ | 58 | 1DP7_P | $\alpha/\beta$ | 76 | 1DTP_A | $\alpha/\beta$ | 190 |
| 1DWM_A | $\alpha/\beta$ | 69 | 1E3Y_A | $\alpha$ | 104 | 1E53_A | $\alpha/\beta$ | 59 | 1EKZ_A | $\alpha/\beta$ | 76 |
| 1ELW_A | $\alpha$ | 117 | 1EM8_D | $\alpha/\beta$ | 112 | 1EZV_G | $\alpha/\beta$ | 81 | 1F15_C | $\alpha/\beta$ | 191 |
| 1F1E_A | $\alpha/\beta$ | 151 | 1F2R_I | $\alpha/\beta$ | 100 | 1F3Y_A | $\alpha/\beta$ | 165 | 1F43_A | $\alpha$ | 61 |
| 1F93_A | $\alpha/\beta$ | 103 | 1F98_A | $\alpha/\beta$ | 125 | 1F9P_A | $\alpha/\beta$ | 81 | 1FAQ_A | $\beta$ | 52 |
| 1FC3_A | $\alpha$ | 116 | 1FCA_A | $\alpha/\beta$ | 55 | 1FEX_A | $\alpha$ | 59 | 1FHT_A | $\alpha/\beta$ | 116 |
| 1FJG_F | $\alpha/\beta$ | 101 | 1FR0_A | $\alpha$ | 125 | 1FRD_A | $\alpha/\beta$ | 98 | 1FSP_A | $\alpha/\beta$ | 124 |
| 1FW9_A | $\alpha/\beta$ | 164 | 1G2R_A | $\alpha/\beta$ | 94 | 1GGS_A | $\alpha/\beta$ | 81 | 1GME_A | $\alpha/\beta$ | 150 |
| 1GPQ_B | $\alpha/\beta$ | 128 | 1GQA_A | $\alpha/\beta$ | 130 | 1GVP_A | $\alpha/\beta$ | 87 | 1GXD_C | $\alpha/\beta$ | 192 |
| 1H9E_A | $\alpha$ | 56 | 1H9F_A | $\alpha$ | 57 | 1HBG_A | $\alpha$ | 147 | 1HBX_E | $\alpha/\beta$ | 89 |
| 1HH8_A | $\alpha/\beta$ | 192 | 1HHV_A | $\alpha/\beta$ | 74 | 1HKQ_A | $\alpha/\beta$ | 125 | 1HKX_E | $\alpha/\beta$ | 143 |
| 1HL6_D | $\alpha/\beta$ | 143 | 1I35_A | $\alpha/\beta$ | 95 | 1I85_A | $\beta$ | 110 | 1IM3_D | $\alpha/\beta$ | 95 |
| 1IOO_A | $\alpha/\beta$ | 196 | 1IS7_K | $\alpha/\beta$ | 85 | 1IUJ_B | $\alpha/\beta$ | 103 | 1IUY_A | $\alpha/\beta$ | 92 |
| 1J8I_A | $\alpha/\beta$ | 93 | 1J9I_A | $\alpha/\beta$ | 68 | 1JEI_A | $\alpha$ | 53 | 1JIW_I | $\alpha/\beta$ | 105 |
| 1JJ2_S | $\alpha/\beta$ | 119 | 1JLI_A | $\alpha$ | 112 | 1JMT_A | $\alpha/\beta$ | 98 | 1JO0_A | $\alpha/\beta$ | 97 |
| 1JOP_A | $\alpha/\beta$ | 140 | 1JPY_Y | $\alpha/\beta$ | 120 | 1JR5_A | $\alpha$ | 90 | 1K1Z_A | $\beta$ | 78 |
| 1K3S_A | $\alpha/\beta$ | 109 | 1K5D_B | $\alpha/\beta$ | 146 | 1K73_1 | $\alpha/\beta$ | 73 | 1KA8_A | $\alpha/\beta$ | 100 |
| 1KN6_A | $\alpha/\beta$ | 73 | 1KOH_D | $\alpha/\beta$ | 172 | 1KPT_A | $\alpha/\beta$ | 105 | 1KQ6_A | $\alpha/\beta$ | 140 |
| 1KSX_A | $\alpha/\beta$ | 144 | 1KX5_D | $\alpha$ | 107 | 1L1D_B | $\alpha/\beta$ | 147 | 1L2P_A | $\alpha$ | 61 |
| 1L3G_A | $\alpha/\beta$ | 123 | 1L6H_A | $\alpha$ | 69 | 1LDD_A | $\alpha/\beta$ | 71 | 1LFU_P | $\alpha$ | 82 |
| 1LNW_C | $\alpha/\beta$ | 139 | 1LR1_B | $\alpha$ | 57 | 1LZW_B | $\alpha/\beta$ | 146 | 1MAI_A | $\alpha/\beta$ | 119 |
| 1MFQ_C | $\alpha$ | 108 | 1MWQ_A | $\alpha/\beta$ | 99 | 1N12_A | $\alpha/\beta$ | 138 | 1N3G_A | $\alpha/\beta$ | 113 |
| 1NF6_F | $\alpha/\beta$ | 171 | 1NGL_A | $\alpha/\beta$ | 179 | 1NKZ_A | $\alpha$ | 53 | 1NOE_A | $\alpha/\beta$ | 86 |
| 1NPB_A | $\alpha/\beta$ | 140 | 1NR3_A | $\alpha/\beta$ | 122 | 1NTV_A | $\alpha/\beta$ | 152 | 1NZE_A | $\alpha$ | 112 |
| 1OFT_A | $\alpha/\beta$ | 119 | 1OJG_A | $\alpha/\beta$ | 136 | 1OOF_A | $\alpha/\beta$ | 124 | 1ORY_A | $\alpha$ | 119 |
| 1OX7_A | $\alpha/\beta$ | 158 | 1OZ9_A | $\alpha/\beta$ | 141 | 1PD6_A | $\beta$ | 94 | 1PGV_A | $\alpha/\beta$ | 167 |
| 1PIH_A | $\alpha/\beta$ | 73 | 1PMS_A | $\alpha/\beta$ | 135 | 1PSR_A | $\alpha/\beta$ | 100 | 1PXW_A | $\alpha/\beta$ | 128 |
| 1PZW_A | $\alpha/\beta$ | 80 | 1QFT_A | $\alpha/\beta$ | 175 | 1QMA_A | $\alpha/\beta$ | 123 | 1QZG_A | $\alpha/\beta$ | 170 |
| 1R5T_A | $\alpha/\beta$ | 141 | 1R6R_A | $\alpha$ | 80 | 1RHX_A | $\alpha/\beta$ | 87 | 1RTU_A | $\alpha/\beta$ | 114 |
| 1RZ3_A | $\alpha/\beta$ | 184 | 1S2D_A | $\alpha/\beta$ | 165 | 1S3J_A | $\alpha/\beta$ | 143 | 1S56_B | $\alpha/\beta$ | 135 |
| 1S7O_C | $\alpha$ | 108 | 1S7Z_A | $\alpha$ | 106 | 1SAU_A | $\alpha/\beta$ | 114 | 1SMP_I | $\alpha/\beta$ | 100 |
| 1SVJ_A | $\alpha/\beta$ | 136 | 1TAF_A | $\alpha$ | 68 | 1TEO_A | $\alpha/\beta$ | 173 | 1TJF_B | $\alpha/\beta$ | 186 |
| 1TLJ_A | $\alpha/\beta$ | 189 | 1TUL_A | $\alpha/\beta$ | 102 | 1TWU_A | $\alpha/\beta$ | 137 | 1TYG_B | $\alpha/\beta$ | 65 |
| 1TZ0_A | $\alpha/\beta$ | 108 | 1U84_A | $\alpha$ | 81 | 1UFB_A | $\alpha$ | 127 | 1UG4_A | $\beta$ | 60 |
| 1UNG_D | $\alpha$ | 149 | 1USL_C | $\alpha/\beta$ | 158 | 1V74_A | $\alpha/\beta$ | 107 | 1VCC_A | $\alpha/\beta$ | 77 |
| 1VCY_A | $\alpha/\beta$ | 193 | 1VHG_A | $\alpha/\beta$ | 185 | 1VKE_E | $\alpha$ | 119 | 1VYX_A | $\alpha/\beta$ | 60 |
| 1W1W_E | $\alpha/\beta$ | 70 | 1WJ8_A | $\alpha$ | 117 | 1WLQ_C | $\alpha/\beta$ | 185 | 1WMH_B | $\alpha/\beta$ | 82 |
| 1XJA_C | $\alpha/\beta$ | 169 | 1Y14_A | $\alpha$ | 133 | 1Y1X_A | $\alpha/\beta$ | 182 | 1YG2_A | $\alpha/\beta$ | 169 |
| 1Z8R_A | $\alpha/\beta$ | 150 | 2A5Y_A | $\alpha$ | 173 | 2A9U_B | $\alpha$ | 127 | 2ACY_A | $\alpha/\beta$ | 98 |
| 2AEN_A | $\alpha/\beta$ | 164 | 2APN_A | $\alpha/\beta$ | 114 | 2AQ0_A | $\alpha$ | 84 | 2AQS_A | $\alpha/\beta$ | 160 |
| 2BSE_A | $\alpha/\beta$ | 107 | 2BWJ_A | $\alpha/\beta$ | 196 | 2BYK_D | $\alpha$ | 92 | 2C2F_A | $\alpha/\beta$ | 178 |
| 2C4W_A | $\alpha/\beta$ | 168 | 2CDP_A | $\alpha/\beta$ | 138 | 2CMX_A | $\alpha/\beta$ | 70 | 2CO3_B | $\alpha/\beta$ | 135 |
| 2CWP_A | $\alpha/\beta$ | 109 | 2CZV_D | $\alpha/\beta$ | 119 | 2D0P_B | $\alpha/\beta$ | 110 | 2EWC_B | $\alpha/\beta$ | 122 |

Continued on next page

Continue to last page

| Protein | Type | Size | Protein | Type | Size | Protein | Type | Size | Protein | Type | Size |
|---|---|---|---|---|---|---|---|---|---|---|---|
| 2F22_B | $\alpha/\beta$ | 143 | 2FA5_B | $\alpha/\beta$ | 142 | 2FKB_C | $\alpha/\beta$ | 167 | 2GBJ_B | $\alpha/\beta$ | 84 |
| 2GJ3_A | $\alpha/\beta$ | 119 | 2H30_A | $\alpha/\beta$ | 151 | 2H8E_A | $\alpha/\beta$ | 120 | 2HI3_A | $\alpha$ | 73 |
| 2HQ7_B | $\alpha/\beta$ | 142 | 2ICT_A | $\alpha$ | 94 | 2J6Z_A | $\alpha$ | 86 | 2JP3_A | $\alpha$ | 67 |
| 2KBW_A | $\alpha$ | 160 | 2L5P_A | $\alpha/\beta$ | 175 | 2L74_A | $\alpha/\beta$ | 125 | 2LKP_A | $\alpha/\beta$ | 119 |
| 2LRB_A | $\alpha/\beta$ | 165 | 2NAZ_A | $\alpha/\beta$ | 109 | 2NCM_A | $\beta$ | 99 | 2NDP_A | $\alpha/\beta$ | 99 |
| 2NS9_B | $\alpha/\beta$ | 152 | 2O70_F | $\alpha$ | 168 | 2ODM_B | $\alpha$ | 83 | 2P7L_A | $\alpha/\beta$ | 125 |
| 2PI2_F | $\alpha/\beta$ | 119 | 2PYB_A | $\alpha$ | 151 | 2Q2H_A | $\alpha/\beta$ | 118 | 2QVG_A | $\alpha/\beta$ | 129 |
| 2QZJ_A | $\alpha/\beta$ | 121 | 2RD5_D | $\alpha/\beta$ | 126 | 2RLD_C | $\alpha$ | 116 | 2UUX_A | $\alpha/\beta$ | 55 |
| 2V85_A | $\alpha/\beta$ | 74 | 2WCW_B | $\alpha/\beta$ | 122 | 2WGP_A | $\alpha/\beta$ | 168 | 2XGY_A | $\alpha$ | 129 |
| 2Z3B_A | $\alpha/\beta$ | 180 | 2ZMZ_B | $\alpha/\beta$ | 79 | 3ALU_A | $\alpha/\beta$ | 157 | 3CAE_A | $\alpha/\beta$ | 132 |
| 3CG4_A | $\alpha/\beta$ | 126 | 3CX5_F | $\alpha$ | 74 | 3CX5_G | $\alpha$ | 126 | 3E6M_E | $\alpha/\beta$ | 147 |
| 3EOD_A | $\alpha/\beta$ | 115 | 3G20_B | $\alpha/\beta$ | 119 | 3GMX_A | $\alpha/\beta$ | 153 | 3I9V_7 | $\alpha/\beta$ | 127 |
| 3IAM_2 | $\alpha/\beta$ | 179 | 3LQV_B | $\alpha/\beta$ | 115 | 3M1N_B | $\alpha/\beta$ | 168 | 3MQK_C | $\beta$ | 75 |
| 3N1G_C | $\alpha/\beta$ | 104 | 3N9U_C | $\alpha/\beta$ | 96 | 3PD2_A | $\alpha/\beta$ | 147 | 3QU3_A | $\alpha/\beta$ | 122 |
| 3SDL_B | $\alpha$ | 97 | 3UE6_E | $\alpha/\beta$ | 138 | 3V1O_A | $\alpha/\beta$ | 165 | 3W1Z_D | $\alpha/\beta$ | 110 |
| 3X0G_A | $\alpha$ | 93 | 3X15_A | $\alpha$ | 87 | 4AIH_A | $\alpha/\beta$ | 139 | 4ASW_C | $\alpha/\beta$ | 81 |
| 4B0M_A | $\alpha/\beta$ | 131 | 4CXT_A | $\alpha/\beta$ | 132 | 4ESB_A | $\alpha/\beta$ | 103 | 4GDK_A | $\alpha/\beta$ | 88 |
| 4GF3_A | $\alpha/\beta$ | 123 | 4GQY_A | $\alpha/\beta$ | 147 | 4I60_A | $\alpha/\beta$ | 128 | 4IOS_A | $\alpha/\beta$ | 100 |
| 4J20_A | $\alpha/\beta$ | 88 | 4JGX_B | $\alpha/\beta$ | 128 | 4KA0_A | $\alpha/\beta$ | 143 | 4LE0_B | $\alpha/\beta$ | 133 |
| 4M75_F | $\alpha/\beta$ | 75 | 4MMG_A | $\alpha/\beta$ | 91 | 4OW1_A | $\alpha/\beta$ | 86 | 4Q2O_A | $\alpha/\beta$ | 92 |
| 4Q2Q_A | $\alpha/\beta$ | 90 | 4R67_0 | $\alpha/\beta$ | 199 | 4RUV_A | $\alpha/\beta$ | 106 | 4UIJ_A | $\alpha/\beta$ | 104 |
| 4V2O_A | $\alpha$ | 78 | 5CJ3_B | $\alpha/\beta$ | 126 | 5E4E_A | $\alpha/\beta$ | 111 | 5EKT_A | $\alpha/\beta$ | 196 |
| 5IAO_A | $\alpha/\beta$ | 171 | 5IZB_A | $\alpha/\beta$ | 89 | 5JTM_A | $\alpha/\beta$ | 155 | 5L38_A | $\alpha/\beta$ | 91 |
| 5L8R_D | $\alpha/\beta$ | 143 | 5O2V_A | $\alpha/\beta$ | 92 | 5O8G_A | $\alpha/\beta$ | 122 | 5T17_A | $\alpha/\beta$ | 85 |
| 5TMF_E | $\alpha/\beta$ | 95 | 5TUV_B | $\alpha/\beta$ | 104 | 5WSE_A | $\alpha/\beta$ | 114 | 6AQ3_B | $\alpha/\beta$ | 171 |

Table S3: Detailed information of 24 CASP13 FM targets.

| Target | Size | Target | Size | Target | Size | Target | Size |
|---|---|---|---|---|---|---|---|
| T0950-D1 | 342 | T0953s1-D1 | 67 | T0953s2-D1 | 44 | T0953s2-D2 | 111 |
| T0953s2-D3 | 93 | T0955-D1 | 41 | T0957s1-D1 | 108 | T0957s2-D1 | 155 |
| T0958-D1 | 77 | T0960-D2 | 84 | T0963-D2 | 82 | T0968s1-D1 | 118 |
| T0968s2-D1 | 115 | T0969-D1 | 354 | T0970-D1 | 85 | T0980s1-D1 | 104 |
| T0990-D1 | 76 | T0990-D2 | 231 | T0990-D3 | 213 | T1005-D1 | 326 |
| T1008-D1 | 77 | T1021s3-D1 | 166 | T1021s3-D2 | 97 | T1022s1-D1 | 156 |

Table S4: Results of the first model of SNfold, Rosetta-dist(500), C-QUARK and SNfold-exploitation on 300 benchmark proteins.

| No. | Protein | SNfold | | Rosetta-dist(500) | | C-QUARK | | SNfold-exploitation | |
|---|---|---|---|---|---|---|---|---|---|
| | | RMSD (Å) | TM-score | RMSD (Å) | TM-score | RMSD (Å) | TM-score | RMSD (Å) | TM-score |
| 1 | 1A3A_C | 6.975 | 0.520 | 5.372 | 0.559 | 3.173 | 0.758 | 11.713 | 0.362 |
| 2 | 1A6L_A | 4.616 | 0.579 | 4.452 | 0.534 | 12.844 | 0.360 | 4.024 | 0.615 |
| 3 | 1A7D_A | 3.195 | 0.764 | 4.122 | 0.709 | 3.311 | 0.760 | 5.305 | 0.675 |
| 4 | 1A91_A | 3.824 | 0.575 | 4.434 | 0.522 | 4.571 | 0.475 | 3.832 | 0.526 |
| 5 | 1ABV_A | 2.508 | 0.768 | 3.043 | 0.720 | 2.909 | 0.758 | 3.900 | 0.610 |
| 6 | 1AHK_A | 7.943 | 0.432 | 5.590 | 0.539 | 5.832 | 0.512 | 10.255 | 0.283 |
| 7 | 1AK6_A | 6.669 | 0.503 | 6.291 | 0.534 | 7.412 | 0.630 | 9.859 | 0.415 |
| 8 | 1AKP_A | 9.506 | 0.401 | 9.128 | 0.402 | 5.637 | 0.588 | 11.732 | 0.264 |
| 9 | 1AP7_A | 4.701 | 0.662 | 4.234 | 0.691 | 4.847 | 0.736 | 4.708 | 0.666 |
| 10 | 1AUU_A | 4.018 | 0.577 | 3.056 | 0.590 | 3.858 | 0.692 | 3.305 | 0.595 |
| 11 | 1AX8_A | 9.597 | 0.523 | 9.711 | 0.396 | 8.133 | 0.428 | 14.116 | 0.450 |
| 12 | 1B4R_A | 4.713 | 0.560 | 3.622 | 0.573 | 2.464 | 0.718 | 4.489 | 0.516 |
| 13 | 1B4U_A | 6.544 | 0.639 | 6.387 | 0.618 | 7.062 | 0.592 | 10.299 | 0.635 |
| 14 | 1BE3_J | 3.745 | 0.560 | 4.599 | 0.500 | 17.216 | 0.271 | 3.759 | 0.523 |
| 15 | 1BGF_A | 3.196 | 0.742 | 3.733 | 0.673 | 4.517 | 0.608 | 3.949 | 0.680 |
| 16 | 1BGY_J | 3.396 | 0.557 | 4.574 | 0.499 | 17.258 | 0.270 | 5.468 | 0.447 |
| 17 | 1BJX_A | 7.551 | 0.640 | 7.148 | 0.624 | 6.439 | 0.741 | 10.941 | 0.429 |
| 18 | 1BUO_A | 4.349 | 0.718 | 5.432 | 0.592 | 9.026 | 0.791 | 8.479 | 0.477 |
| 19 | 1C03_A | 12.790 | 0.649 | 12.841 | 0.556 | 7.876 | 0.679 | 15.327 | 0.579 |
| 20 | 1C41_A | 5.451 | 0.657 | 11.348 | 0.360 | 4.722 | 0.712 | 8.744 | 0.441 |
| 21 | 1C9F_A | 3.033 | 0.668 | 3.812 | 0.623 | 6.679 | 0.601 | 5.349 | 0.565 |
| 22 | 1CDB_A | 6.742 | 0.519 | 6.039 | 0.464 | 4.933 | 0.665 | 8.451 | 0.350 |
| 23 | 1CF7_B | 3.537 | 0.694 | 2.952 | 0.673 | 2.823 | 0.684 | 3.122 | 0.696 |
| 24 | 1CTO_A | 8.696 | 0.371 | 9.022 | 0.283 | 5.504 | 0.627 | 9.892 | 0.338 |
| 25 | 1CXZ_B | 1.774 | 0.852 | 1.827 | 0.845 | 2.962 | 0.797 | 2.168 | 0.829 |
| 26 | 1D6T_A | 5.579 | 0.592 | 5.118 | 0.587 | 3.645 | 0.705 | 5.944 | 0.559 |
| 27 | 1D8B_A | 3.254 | 0.666 | 3.392 | 0.591 | 3.140 | 0.659 | 3.228 | 0.644 |
| 28 | 1DBF_A | 8.229 | 0.604 | 6.772 | 0.507 | 5.477 | 0.639 | 13.787 | 0.359 |
| 29 | 1DCF_A | 5.228 | 0.725 | 3.968 | 0.751 | 3.996 | 0.823 | 6.174 | 0.532 |
| 30 | 1DL6_A | 13.678 | 0.391 | 12.768 | 0.401 | 12.674 | 0.369 | 13.566 | 0.374 |
| 31 | 1DP7_P | 3.390 | 0.711 | 3.031 | 0.649 | 3.378 | 0.744 | 2.415 | 0.720 |
| 32 | 1DTP_A | 15.770 | 0.238 | 19.327 | 0.185 | 17.420 | 0.223 | 15.842 | 0.214 |
| 33 | 1DWM_A | 4.189 | 0.643 | 3.855 | 0.611 | 5.133 | 0.643 | 5.034 | 0.637 |
| 34 | 1E3Y_A | 5.080 | 0.724 | 8.088 | 0.652 | 5.789 | 0.685 | 4.933 | 0.670 |
| 35 | 1E53_A | 3.230 | 0.592 | 4.006 | 0.454 | 3.915 | 0.516 | 3.913 | 0.571 |
| 36 | 1EKZ_A | 5.347 | 0.678 | 4.533 | 0.654 | 5.577 | 0.690 | 5.523 | 0.653 |
| 37 | 1ELW_A | 1.252 | 0.920 | 1.365 | 0.907 | 1.661 | 0.870 | 1.368 | 0.907 |
| 38 | 1EM8_D | 7.097 | 0.491 | 6.138 | 0.491 | 6.268 | 0.568 | 7.428 | 0.338 |
| 39 | 1EZV_G | 12.593 | 0.417 | 10.349 | 0.412 | 25.700 | 0.324 | 11.061 | 0.422 |
| 40 | 1F15_C | 21.426 | 0.204 | 21.117 | 0.221 | 17.510 | 0.215 | 21.091 | 0.199 |
| 41 | 1F1E_A | 11.397 | 0.489 | 13.306 | 0.468 | 7.676 | 0.476 | 10.652 | 0.497 |
| 42 | 1F2R_I | 5.486 | 0.635 | 4.582 | 0.593 | 9.012 | 0.643 | 6.002 | 0.561 |
| 43 | 1F3Y_A | 15.273 | 0.329 | 13.833 | 0.305 | 9.523 | 0.513 | 13.171 | 0.298 |
| 44 | 1F43_A | 4.701 | 0.583 | 5.980 | 0.563 | 8.033 | 0.559 | 7.388 | 0.563 |
| 45 | 1F93_A | 2.262 | 0.794 | 2.812 | 0.716 | 1.816 | 0.860 | 3.284 | 0.664 |
| 46 | 1F98_A | 9.222 | 0.575 | 8.542 | 0.593 | 3.671 | 0.658 | 11.073 | 0.490 |
| 47 | 1F9P_A | 7.281 | 0.616 | 5.249 | 0.576 | 12.778 | 0.611 | 7.802 | 0.566 |
| 48 | 1FAQ_A | 3.476 | 0.551 | 4.688 | 0.446 | 4.600 | 0.402 | 5.216 | 0.484 |

Continued on next page

Continue to last page

| No. | Protein | SNfold | | Rosetta-dist(500) | | C-QUARK | | SNfold-exploitation | |
|---|---|---|---|---|---|---|---|---|---|
| | | RMSD (Å) | TM-score | RMSD (Å) | TM-score | RMSD (Å) | TM-score | RMSD (Å) | TM-score |
| 49 | 1FC3_A | 2.945 | 0.778 | 3.369 | 0.680 | 3.194 | 0.772 | 5.173 | 0.597 |
| 50 | 1FCA_A | 1.184 | 0.821 | 1.860 | 0.699 | 2.174 | 0.675 | 1.594 | 0.745 |
| 51 | 1FEX_A | 2.648 | 0.669 | 2.867 | 0.614 | 2.717 | 0.587 | 2.546 | 0.651 |
| 52 | 1FHT_A | 13.350 | 0.559 | 11.152 | 0.518 | 10.474 | 0.663 | 11.743 | 0.360 |
| 53 | 1FJG_F | 6.493 | 0.660 | 5.292 | 0.604 | 3.591 | 0.809 | 7.213 | 0.570 |
| 54 | 1FR0_A | 3.872 | 0.756 | 3.198 | 0.721 | 4.365 | 0.699 | 4.156 | 0.621 |
| 55 | 1FRD_A | 4.986 | 0.639 | 4.848 | 0.535 | 3.646 | 0.704 | 6.804 | 0.476 |
| 56 | 1FSP_A | 3.295 | 0.830 | 3.697 | 0.735 | 3.256 | 0.843 | 3.869 | 0.706 |
| 57 | 1FW9_A | 12.159 | 0.476 | 10.316 | 0.447 | 8.315 | 0.607 | 15.631 | 0.241 |
| 58 | 1G2R_A | 3.483 | 0.756 | 2.979 | 0.697 | 2.382 | 0.771 | 3.594 | 0.675 |
| 59 | 1GGS_A | 9.258 | 0.707 | 8.912 | 0.694 | 10.681 | 0.659 | 7.278 | 0.693 |
| 60 | 1GME_A | 15.285 | 0.419 | 15.512 | 0.338 | 14.425 | 0.478 | 13.689 | 0.358 |
| 61 | 1GPQ_B | 7.349 | 0.545 | 6.744 | 0.546 | 5.142 | 0.664 | 12.381 | 0.352 |
| 62 | 1GQA_A | 5.489 | 0.735 | 6.746 | 0.722 | 3.967 | 0.756 | 9.307 | 0.549 |
| 63 | 1GVP_A | 4.603 | 0.639 | 4.733 | 0.542 | 5.369 | 0.595 | 5.679 | 0.551 |
| 64 | 1GXD_C | 12.333 | 0.383 | 15.770 | 0.253 | 11.630 | 0.427 | 21.795 | 0.233 |
| 65 | 1H9E_A | 5.787 | 0.491 | 6.061 | 0.510 | 7.556 | 0.521 | 5.681 | 0.489 |
| 66 | 1H9F_A | 3.916 | 0.553 | 3.998 | 0.542 | 4.161 | 0.595 | 3.773 | 0.544 |
| 67 | 1HBG_A | 3.342 | 0.777 | 2.796 | 0.799 | 2.205 | 0.840 | 7.944 | 0.540 |
| 68 | 1HBX_E | 7.981 | 0.599 | 8.827 | 0.545 | 12.568 | 0.414 | 10.779 | 0.533 |
| 69 | 1HH8_A | 12.938 | 0.740 | 16.931 | 0.677 | 15.207 | 0.565 | 14.669 | 0.689 |
| 70 | 1HHV_A | 6.926 | 0.575 | 6.651 | 0.507 | 11.293 | 0.628 | 10.514 | 0.406 |
| 71 | 1HKQ_A | 4.205 | 0.662 | 3.649 | 0.637 | 5.191 | 0.610 | 6.270 | 0.479 |
| 72 | 1HKX_E | 7.857 | 0.483 | 10.216 | 0.509 | 5.412 | 0.657 | 9.929 | 0.361 |
| 73 | 1HL6_D | 7.946 | 0.494 | 9.665 | 0.363 | 5.444 | 0.540 | 11.876 | 0.346 |
| 74 | 1I35_A | 4.816 | 0.603 | 5.090 | 0.525 | 5.334 | 0.439 | 5.323 | 0.568 |
| 75 | 1I85_A | 6.481 | 0.553 | 5.913 | 0.528 | 3.856 | 0.605 | 12.064 | 0.495 |
| 76 | 1IM3_D | 13.891 | 0.302 | 12.130 | 0.303 | 12.444 | 0.278 | 12.727 | 0.255 |
| 77 | 1IOO_A | 11.599 | 0.368 | 11.754 | 0.288 | 7.901 | 0.530 | 13.301 | 0.350 |
| 78 | 1IS7_K | 7.289 | 0.612 | 9.255 | 0.378 | 7.493 | 0.484 | 8.624 | 0.445 |
| 79 | 1IUJ_B | 3.803 | 0.772 | 3.653 | 0.687 | 4.588 | 0.706 | 3.620 | 0.724 |
| 80 | 1IUY_A | 8.803 | 0.621 | 9.633 | 0.480 | 11.984 | 0.677 | 12.160 | 0.282 |
| 81 | 1J8I_A | 14.701 | 0.567 | 15.165 | 0.554 | 19.233 | 0.568 | 17.580 | 0.577 |
| 82 | 1J9I_A | 6.027 | 0.590 | 6.425 | 0.533 | 4.863 | 0.632 | 5.819 | 0.552 |
| 83 | 1JEI_A | 6.804 | 0.597 | 6.392 | 0.600 | 8.138 | 0.610 | 7.369 | 0.589 |
| 84 | 1JIW_I | 7.660 | 0.574 | 5.865 | 0.489 | 4.146 | 0.662 | 10.161 | 0.345 |
| 85 | 1JJ2_S | 5.464 | 0.628 | 5.522 | 0.540 | 10.464 | 0.437 | 7.641 | 0.471 |
| 86 | 1JLI_A | 12.284 | 0.384 | 17.338 | 0.237 | 10.629 | 0.336 | 14.917 | 0.345 |
| 87 | 1JMT_A | 5.039 | 0.627 | 5.429 | 0.584 | 5.294 | 0.701 | 8.739 | 0.587 |
| 88 | 1JO0_A | 1.935 | 0.830 | 2.439 | 0.742 | 2.618 | 0.806 | 3.622 | 0.584 |
| 89 | 1JOP_A | 13.261 | 0.451 | 10.335 | 0.353 | 4.183 | 0.628 | 9.595 | 0.329 |
| 90 | 1JPY_Y | 21.326 | 0.354 | 14.629 | 0.311 | 12.871 | 0.472 | 12.971 | 0.343 |
| 91 | 1JR5_A | 8.602 | 0.574 | 8.483 | 0.444 | 8.521 | 0.345 | 9.158 | 0.428 |
| 92 | 1K1Z_A | 9.573 | 0.563 | 5.972 | 0.481 | 5.137 | 0.602 | 8.789 | 0.465 |
| 93 | 1K3S_A | 8.097 | 0.502 | 4.967 | 0.538 | 12.462 | 0.299 | 12.489 | 0.333 |
| 94 | 1K5D_B | 9.816 | 0.492 | 8.661 | 0.476 | 9.773 | 0.618 | 13.299 | 0.343 |
| 95 | 1K73_1 | 3.853 | 0.670 | 3.042 | 0.616 | 6.156 | 0.447 | 4.586 | 0.575 |
| 96 | 1KA8_A | 5.848 | 0.611 | 6.976 | 0.546 | 6.478 | 0.470 | 7.230 | 0.523 |
| 97 | 1KN6_A | 3.864 | 0.586 | 3.586 | 0.554 | 4.210 | 0.591 | 4.656 | 0.455 |
| 98 | 1KOH_D | 8.006 | 0.649 | 9.106 | 0.703 | 9.915 | 0.544 | 7.011 | 0.641 |

Continued on next page

Continue to last page

| No. | Protein | SNfold | | Rosetta-dist(500) | | C-QUARK | | SNfold-exploitation | |
|---|---|---|---|---|---|---|---|---|---|
| | | RMSD (Å) | TM-score | RMSD (Å) | TM-score | RMSD (Å) | TM-score | RMSD (Å) | TM-score |
| 99 | 1KPT_A | 5.901 | 0.458 | 11.011 | 0.341 | 4.381 | 0.619 | 11.576 | 0.352 |
| 100 | 1KQ6_A | 7.516 | 0.587 | 9.748 | 0.527 | 4.753 | 0.722 | 11.435 | 0.518 |
| 101 | 1KSX_A | 5.780 | 0.540 | 5.468 | 0.536 | 3.268 | 0.730 | 10.675 | 0.395 |
| 102 | 1KX5_D | 6.383 | 0.814 | 5.186 | 0.807 | 9.738 | 0.477 | 8.366 | 0.795 |
| 103 | 1L1D_B | 8.146 | 0.376 | 10.181 | 0.359 | 4.401 | 0.655 | 14.066 | 0.259 |
| 104 | 1L2P_A | 0.622 | 0.949 | 0.890 | 0.909 | 25.047 | 0.458 | 0.630 | 0.949 |
| 105 | 1L3G_A | 7.313 | 0.537 | 11.648 | 0.555 | 8.411 | 0.557 | 13.398 | 0.469 |
| 106 | 1L6H_A | 4.350 | 0.512 | 6.755 | 0.495 | 4.849 | 0.413 | 4.658 | 0.442 |
| 107 | 1LDD_A | 2.208 | 0.790 | 2.598 | 0.702 | 2.291 | 0.708 | 3.427 | 0.614 |
| 108 | 1LFU_P | 11.651 | 0.591 | 11.794 | 0.589 | 11.208 | 0.567 | 11.926 | 0.566 |
| 109 | 1LNW_C | 3.307 | 0.854 | 3.856 | 0.758 | 10.552 | 0.663 | 2.901 | 0.822 |
| 110 | 1LR1_B | 3.867 | 0.536 | 4.042 | 0.515 | 5.365 | 0.434 | 3.783 | 0.518 |
| 111 | 1LZW_B | 4.827 | 0.752 | 6.402 | 0.682 | 3.112 | 0.815 | 5.838 | 0.598 |
| 112 | 1MAI_A | 6.133 | 0.530 | 4.899 | 0.518 | 3.728 | 0.705 | 13.475 | 0.266 |
| 113 | 1MFQ_C | 8.425 | 0.627 | 9.114 | 0.566 | 9.833 | 0.583 | 9.383 | 0.551 |
| 114 | 1MWQ_A | 4.756 | 0.645 | 3.657 | 0.607 | 6.255 | 0.576 | 8.502 | 0.407 |
| 115 | 1N12_A | 11.512 | 0.353 | 13.230 | 0.245 | 5.977 | 0.489 | 11.861 | 0.230 |
| 116 | 1N3G_A | 6.432 | 0.668 | 6.680 | 0.629 | 5.055 | 0.753 | 10.869 | 0.406 |
| 117 | 1NF6_F | 8.008 | 0.749 | 12.030 | 0.658 | 8.957 | 0.815 | 8.576 | 0.511 |
| 118 | 1NGL_A | 9.804 | 0.493 | 8.586 | 0.449 | 8.645 | 0.602 | 10.144 | 0.373 |
| 119 | 1NKZ_A | 6.301 | 0.612 | 5.405 | 0.559 | 14.598 | 0.417 | 5.832 | 0.562 |
| 120 | 1NOE_A | 7.189 | 0.392 | 5.288 | 0.430 | 5.191 | 0.515 | 6.876 | 0.336 |
| 121 | 1NPB_A | 5.543 | 0.616 | 8.014 | 0.589 | 9.131 | 0.485 | 8.412 | 0.559 |
| 122 | 1NR3_A | 13.166 | 0.337 | 13.329 | 0.268 | 12.009 | 0.272 | 20.741 | 0.312 |
| 123 | 1NTV_A | 7.707 | 0.503 | 6.151 | 0.477 | 4.654 | 0.695 | 12.009 | 0.280 |
| 124 | 1NZE_A | 2.266 | 0.865 | 4.292 | 0.813 | 2.334 | 0.869 | 3.013 | 0.825 |
| 125 | 1OFT_A | 3.309 | 0.702 | 4.427 | 0.575 | 2.566 | 0.776 | 3.238 | 0.681 |
| 126 | 1OJG_A | 6.943 | 0.578 | 5.267 | 0.522 | 3.861 | 0.633 | 7.491 | 0.489 |
| 127 | 1OOF_A | 3.315 | 0.737 | 6.209 | 0.672 | 2.756 | 0.756 | 2.742 | 0.749 |
| 128 | 1ORY_A | 3.178 | 0.775 | 3.902 | 0.678 | 3.493 | 0.731 | 3.771 | 0.685 |
| 129 | 1OX7_A | 7.527 | 0.552 | 5.235 | 0.565 | 3.238 | 0.792 | 13.081 | 0.321 |
| 130 | 1OZ9_A | 4.839 | 0.649 | 8.309 | 0.563 | 2.732 | 0.807 | 10.876 | 0.544 |
| 131 | 1PD6_A | 6.049 | 0.539 | 4.556 | 0.644 | 4.311 | 0.662 | 9.517 | 0.364 |
| 132 | 1PGV_A | 4.862 | 0.697 | 6.521 | 0.618 | 3.006 | 0.788 | 4.101 | 0.686 |
| 133 | 1PIH_A | 8.365 | 0.428 | 6.972 | 0.403 | 4.666 | 0.519 | 10.028 | 0.216 |
| 134 | 1PMS_A | 8.434 | 0.510 | 12.109 | 0.492 | 8.256 | 0.614 | 9.512 | 0.494 |
| 135 | 1PSR_A | 4.424 | 0.708 | 3.861 | 0.635 | 5.738 | 0.660 | 3.615 | 0.657 |
| 136 | 1PXW_A | 5.830 | 0.530 | 5.791 | 0.501 | 3.716 | 0.757 | 11.110 | 0.359 |
| 137 | 1PZW_A | 6.264 | 0.581 | 4.518 | 0.472 | 11.697 | 0.523 | 5.491 | 0.558 |
| 138 | 1QFT_A | 7.906 | 0.464 | 14.721 | 0.278 | 4.686 | 0.689 | 10.264 | 0.355 |
| 139 | 1QMA_A | 5.425 | 0.591 | 4.714 | 0.562 | 2.255 | 0.893 | 7.107 | 0.470 |
| 140 | 1QZG_A | 12.037 | 0.397 | 9.043 | 0.419 | 5.186 | 0.687 | 9.463 | 0.463 |
| 141 | 1R5T_A | 6.763 | 0.557 | 5.422 | 0.586 | 3.879 | 0.739 | 10.376 | 0.409 |
| 142 | 1R6R_A | 8.144 | 0.535 | 8.636 | 0.556 | 8.710 | 0.385 | 8.247 | 0.524 |
| 143 | 1RHX_A | 5.024 | 0.509 | 4.625 | 0.486 | 4.610 | 0.568 | 5.226 | 0.537 |
| 144 | 1RTU_A | 6.513 | 0.555 | 6.980 | 0.529 | 4.460 | 0.683 | 11.936 | 0.490 |
| 145 | 1RZ3_A | 11.552 | 0.489 | 8.849 | 0.511 | 7.512 | 0.604 | 12.577 | 0.481 |
| 146 | 1S2D_A | 8.857 | 0.534 | 7.615 | 0.542 | 4.688 | 0.721 | 14.379 | 0.307 |
| 147 | 1S3J_A | 3.317 | 0.782 | 2.889 | 0.753 | 8.108 | 0.752 | 6.641 | 0.605 |
| 148 | 1S56_B | 7.297 | 0.653 | 7.271 | 0.670 | 8.081 | 0.724 | 9.059 | 0.506 |

Continued on next page

Continue to last page

| No. | Protein | SNfold | | Rosetta-dist(500) | | C-QUARK | | SNfold-exploitation | |
|---|---|---|---|---|---|---|---|---|---|
| | | RMSD (Å) | TM-score | RMSD (Å) | TM-score | RMSD (Å) | TM-score | RMSD (Å) | TM-score |
| 149 | 1S7O_C | 10.538 | 0.585 | 11.383 | 0.557 | 5.069 | 0.679 | 12.115 | 0.592 |
| 150 | 1S7Z_A | 8.135 | 0.582 | 7.840 | 0.508 | 4.513 | 0.686 | 7.760 | 0.502 |
| 151 | 1SAU_A | 3.761 | 0.675 | 4.299 | 0.643 | 3.285 | 0.672 | 8.317 | 0.556 |
| 152 | 1SMP_I | 9.162 | 0.563 | 5.725 | 0.523 | 3.016 | 0.694 | 5.296 | 0.561 |
| 153 | 1SVJ_A | 5.523 | 0.604 | 6.585 | 0.472 | 3.642 | 0.757 | 9.912 | 0.444 |
| 154 | 1TAF_A | 1.231 | 0.887 | 1.165 | 0.894 | 6.552 | 0.513 | 1.233 | 0.894 |
| 155 | 1TEO_A | 11.633 | 0.555 | 11.354 | 0.452 | 7.682 | 0.647 | 11.184 | 0.498 |
| 156 | 1TJF_B | 6.138 | 0.731 | 5.444 | 0.722 | 4.013 | 0.772 | 6.920 | 0.630 |
| 157 | 1TLJ_A | 10.932 | 0.436 | 7.364 | 0.492 | 10.619 | 0.470 | 12.773 | 0.384 |
| 158 | 1TUL_A | 9.036 | 0.400 | 8.478 | 0.341 | 8.568 | 0.460 | 9.651 | 0.312 |
| 159 | 1TWU_A | 6.270 | 0.546 | 3.953 | 0.689 | 5.730 | 0.606 | 5.249 | 0.582 |
| 160 | 1TYG_B | 2.141 | 0.733 | 2.683 | 0.672 | 3.162 | 0.803 | 2.610 | 0.679 |
| 161 | 1TZ0_A | 6.321 | 0.550 | 5.661 | 0.489 | 5.110 | 0.650 | 6.645 | 0.516 |
| 162 | 1U84_A | 1.738 | 0.843 | 1.934 | 0.825 | 2.003 | 0.800 | 2.990 | 0.799 |
| 163 | 1UFB_A | 4.626 | 0.732 | 6.842 | 0.616 | 2.744 | 0.801 | 12.355 | 0.599 |
| 164 | 1UG4_A | 5.012 | 0.447 | 5.408 | 0.379 | 4.228 | 0.440 | 5.848 | 0.619 |
| 165 | 1UNG_D | 4.373 | 0.646 | 4.554 | 0.640 | 7.950 | 0.503 | 9.409 | 0.405 |
| 166 | 1USL_C | 5.482 | 0.624 | 5.226 | 0.626 | 8.383 | 0.775 | 8.156 | 0.440 |
| 167 | 1V74_A | 6.163 | 0.645 | 4.881 | 0.623 | 4.933 | 0.596 | 5.756 | 0.555 |
| 168 | 1VCC_A | 4.500 | 0.542 | 4.216 | 0.563 | 4.989 | 0.552 | 7.596 | 0.435 |
| 169 | 1VCY_A | 11.622 | 0.395 | 13.509 | 0.302 | 14.468 | 0.322 | 15.041 | 0.271 |
| 170 | 1VHG_A | 10.620 | 0.426 | 10.511 | 0.338 | 14.456 | 0.611 | 13.677 | 0.360 |
| 171 | 1VKE_E | 6.109 | 0.727 | 5.661 | 0.662 | 11.481 | 0.642 | 5.072 | 0.699 |
| 172 | 1VYX_A | 7.475 | 0.442 | 5.673 | 0.355 | 4.922 | 0.433 | 7.774 | 0.358 |
| 173 | 1W1W_E | 2.518 | 0.743 | 2.564 | 0.712 | 1.920 | 0.813 | 2.275 | 0.710 |
| 174 | 1WJ8_A | 2.328 | 0.812 | 4.560 | 0.758 | 1.958 | 0.839 | 6.953 | 0.583 |
| 175 | 1WLQ_C | 10.450 | 0.501 | 9.863 | 0.460 | 8.298 | 0.576 | 14.800 | 0.313 |
| 176 | 1WMH_B | 3.233 | 0.640 | 3.626 | 0.561 | 2.257 | 0.733 | 4.566 | 0.494 |
| 177 | 1XJA_C | 6.084 | 0.665 | 4.951 | 0.593 | 4.940 | 0.691 | 8.220 | 0.515 |
| 178 | 1Y14_A | 9.028 | 0.509 | 9.016 | 0.432 | 6.655 | 0.508 | 12.798 | 0.432 |
| 179 | 1Y1X_A | 8.824 | 0.638 | 6.787 | 0.620 | 4.985 | 0.629 | 8.882 | 0.620 |
| 180 | 1YG2_A | 5.293 | 0.558 | 12.790 | 0.410 | 17.175 | 0.438 | 10.049 | 0.458 |
| 181 | 1Z8R_A | 13.468 | 0.312 | 13.007 | 0.297 | 14.707 | 0.297 | 16.452 | 0.253 |
| 182 | 2A5Y_A | 4.541 | 0.700 | 4.907 | 0.645 | 12.280 | 0.623 | 15.954 | 0.330 |
| 183 | 2A9U_B | 10.806 | 0.757 | 9.216 | 0.754 | 11.197 | 0.719 | 10.408 | 0.699 |
| 184 | 2ACY_A | 3.470 | 0.670 | 3.653 | 0.626 | 2.461 | 0.859 | 4.981 | 0.559 |
| 185 | 2AEN_A | 13.133 | 0.381 | 13.594 | 0.287 | 14.422 | 0.261 | 16.969 | 0.280 |
| 186 | 2APN_A | 5.825 | 0.538 | 6.894 | 0.492 | 6.114 | 0.666 | 7.586 | 0.490 |
| 187 | 2AQ0_A | 7.690 | 0.658 | 7.237 | 0.643 | 8.483 | 0.630 | 7.380 | 0.645 |
| 188 | 2AQS_A | 10.263 | 0.411 | 11.967 | 0.324 | 5.865 | 0.607 | 11.221 | 0.330 |
| 189 | 2BSE_A | 10.783 | 0.441 | 12.810 | 0.334 | 10.096 | 0.306 | 12.885 | 0.402 |
| 190 | 2BWJ_A | 7.745 | 0.556 | 7.288 | 0.526 | 4.542 | 0.718 | 9.628 | 0.439 |
| 191 | 2BYK_D | 1.764 | 0.870 | 1.824 | 0.852 | 4.634 | 0.570 | 2.274 | 0.795 |
| 192 | 2C2F_A | 9.078 | 0.740 | 6.807 | 0.693 | 5.833 | 0.802 | 8.534 | 0.558 |
| 193 | 2C4W_A | 7.311 | 0.536 | 6.281 | 0.505 | 4.602 | 0.723 | 13.873 | 0.338 |
| 194 | 2CDP_A | 10.584 | 0.371 | 7.770 | 0.446 | 5.093 | 0.630 | 11.650 | 0.316 |
| 195 | 2CMX_A | 2.404 | 0.719 | 3.187 | 0.649 | 3.174 | 0.596 | 4.209 | 0.644 |
| 196 | 2CO3_B | 10.913 | 0.332 | 10.631 | 0.292 | 6.943 | 0.499 | 12.351 | 0.271 |
| 197 | 2CWP_A | 6.367 | 0.562 | 6.130 | 0.478 | 5.070 | 0.725 | 13.836 | 0.240 |
| 198 | 2CZV_D | 7.102 | 0.619 | 5.488 | 0.620 | 5.438 | 0.670 | 10.434 | 0.415 |

Continued on next page

Continue to last page

| No. | Protein | SNfold | | Rosetta-dist(500) | | C-QUARK | | SNfold-exploitation | |
|---|---|---|---|---|---|---|---|---|---|
| | | RMSD (Å) | TM-score | RMSD (Å) | TM-score | RMSD (Å) | TM-score | RMSD (Å) | TM-score |
| 199 | 2D0P_B | 4.506 | 0.649 | 8.774 | 0.638 | 3.193 | 0.747 | 5.835 | 0.509 |
| 200 | 2EWC_B | 5.272 | 0.639 | 7.740 | 0.637 | 3.899 | 0.746 | 7.066 | 0.549 |
| 201 | 2F22_B | 5.798 | 0.565 | 5.672 | 0.563 | 4.664 | 0.737 | 12.410 | 0.391 |
| 202 | 2FA5_B | 5.535 | 0.717 | 4.535 | 0.708 | 9.864 | 0.625 | 5.159 | 0.713 |
| 203 | 2FKB_C | 8.642 | 0.445 | 9.184 | 0.425 | 4.506 | 0.635 | 15.029 | 0.336 |
| 204 | 2GBJ_B | 3.624 | 0.725 | 4.095 | 0.627 | 3.338 | 0.779 | 4.264 | 0.549 |
| 205 | 2GJ3_A | 4.419 | 0.664 | 4.789 | 0.633 | 3.181 | 0.833 | 4.539 | 0.650 |
| 206 | 2H30_A | 8.887 | 0.574 | 6.956 | 0.549 | 6.320 | 0.713 | 12.496 | 0.503 |
| 207 | 2H8E_A | 5.817 | 0.514 | 5.502 | 0.506 | 3.904 | 0.628 | 9.368 | 0.357 |
| 208 | 2HI3_A | 6.801 | 0.633 | 5.657 | 0.627 | 4.945 | 0.695 | 5.481 | 0.618 |
| 209 | 2HQ7_B | 5.529 | 0.529 | 5.705 | 0.535 | 4.120 | 0.712 | 9.267 | 0.468 |
| 210 | 2ICT_A | 3.992 | 0.763 | 5.248 | 0.759 | 5.235 | 0.750 | 5.499 | 0.746 |
| 211 | 2J6Z_A | 1.928 | 0.825 | 2.517 | 0.733 | 5.186 | 0.666 | 1.920 | 0.821 |
| 212 | 2JP3_A | 8.769 | 0.382 | 6.035 | 0.361 | 20.702 | 0.257 | 7.036 | 0.374 |
| 213 | 2KBW_A | 4.573 | 0.694 | 4.283 | 0.715 | 4.847 | 0.693 | 8.551 | 0.550 |
| 214 | 2L5P_A | 4.882 | 0.679 | 6.606 | 0.515 | 4.580 | 0.658 | 7.986 | 0.531 |
| 215 | 2L74_A | 5.203 | 0.705 | 5.348 | 0.546 | 9.059 | 0.646 | 7.409 | 0.487 |
| 216 | 2LKP_A | 7.995 | 0.649 | 9.587 | 0.607 | 11.580 | 0.561 | 11.520 | 0.592 |
| 217 | 2LRB_A | 7.264 | 0.485 | 14.704 | 0.259 | 4.896 | 0.679 | 15.978 | 0.282 |
| 218 | 2NAZ_A | 4.904 | 0.670 | 4.086 | 0.649 | 4.189 | 0.747 | 5.302 | 0.567 |
| 219 | 2NCM_A | 4.354 | 0.610 | 4.241 | 0.594 | 1.667 | 0.861 | 9.429 | 0.407 |
| 220 | 2NDP_A | 6.248 | 0.527 | 7.572 | 0.478 | 11.508 | 0.416 | 7.743 | 0.407 |
| 221 | 2NS9_B | 5.120 | 0.697 | 6.513 | 0.525 | 4.875 | 0.718 | 13.478 | 0.299 |
| 222 | 2O70_F | 8.006 | 0.606 | 5.940 | 0.551 | 2.869 | 0.782 | 7.688 | 0.557 |
| 223 | 2ODM_B | 1.793 | 0.874 | 1.845 | 0.852 | 2.935 | 0.682 | 2.032 | 0.821 |
| 224 | 2P7L_A | 4.850 | 0.627 | 4.230 | 0.637 | 6.196 | 0.654 | 6.202 | 0.514 |
| 225 | 2PI2_F | 6.681 | 0.670 | 5.180 | 0.539 | 4.586 | 0.743 | 10.548 | 0.362 |
| 226 | 2PYB_A | 3.936 | 0.732 | 4.509 | 0.690 | 3.205 | 0.858 | 5.491 | 0.672 |
| 227 | 2Q2H_A | 8.784 | 0.456 | 12.359 | 0.277 | 7.449 | 0.664 | 16.728 | 0.242 |
| 228 | 2QVG_A | 3.535 | 0.752 | 3.635 | 0.713 | 2.757 | 0.810 | 4.950 | 0.628 |
| 229 | 2QZJ_A | 2.345 | 0.819 | 4.314 | 0.751 | 1.253 | 0.923 | 4.781 | 0.691 |
| 230 | 2RD5_D | 8.094 | 0.500 | 10.560 | 0.438 | 12.723 | 0.530 | 15.101 | 0.468 |
| 231 | 2RLD_C | 2.480 | 0.870 | 2.852 | 0.783 | 3.230 | 0.825 | 3.653 | 0.852 |
| 232 | 2UUX_A | 4.346 | 0.525 | 4.714 | 0.387 | 6.489 | 0.355 | 6.105 | 0.353 |
| 233 | 2V85_A | 7.826 | 0.480 | 7.533 | 0.402 | 11.657 | 0.275 | 8.650 | 0.432 |
| 234 | 2WCW_B | 4.304 | 0.682 | 4.378 | 0.610 | 3.790 | 0.749 | 5.501 | 0.570 |
| 235 | 2WGP_A | 6.047 | 0.667 | 4.982 | 0.662 | 4.865 | 0.835 | 7.839 | 0.637 |
| 236 | 2XGY_A | 10.190 | 0.617 | 7.012 | 0.569 | 9.062 | 0.436 | 14.321 | 0.630 |
| 237 | 2Z3B_A | 9.897 | 0.484 | 14.397 | 0.209 | 4.480 | 0.661 | 13.454 | 0.319 |
| 238 | 2ZMZ_B | 4.421 | 0.629 | 4.953 | 0.547 | 9.470 | 0.456 | 5.138 | 0.546 |
| 239 | 3ALU_A | 12.737 | 0.371 | 8.816 | 0.340 | 5.202 | 0.630 | 12.607 | 0.279 |
| 240 | 3CAE_A | 24.345 | 0.452 | 23.747 | 0.421 | 24.815 | 0.471 | 24.615 | 0.352 |
| 241 | 3CG4_A | 3.027 | 0.776 | 3.718 | 0.743 | 2.755 | 0.845 | 3.894 | 0.683 |
| 242 | 3CX5_F | 3.034 | 0.715 | 3.631 | 0.595 | 3.764 | 0.709 | 4.303 | 0.625 |
| 243 | 3CX5_G | 4.109 | 0.642 | 6.647 | 0.459 | 8.960 | 0.544 | 11.133 | 0.468 |
| 244 | 3E6M_E | 4.001 | 0.712 | 9.008 | 0.625 | 5.497 | 0.618 | 6.038 | 0.621 |
| 245 | 3EOD_A | 4.011 | 0.795 | 4.442 | 0.721 | 2.423 | 0.795 | 5.640 | 0.716 |
| 246 | 3G20_B | 8.126 | 0.527 | 8.226 | 0.503 | 8.935 | 0.618 | 6.310 | 0.573 |
| 247 | 3GMX_A | 10.612 | 0.376 | 8.089 | 0.357 | 12.484 | 0.340 | 12.320 | 0.309 |
| 248 | 3I9V_7 | 7.091 | 0.479 | 7.500 | 0.431 | 10.474 | 0.423 | 10.284 | 0.435 |

Continued on next page

Continue to last page

| No. | Protein | SNfold | | Rosetta-dist(500) | | C-QUARK | | SNfold-exploitation | |
|---|---|---|---|---|---|---|---|---|---|
| | | RMSD (Å) | TM-score | RMSD (Å) | TM-score | RMSD (Å) | TM-score | RMSD (Å) | TM-score |
| 249 | 3IAM_2 | 5.936 | 0.617 | 6.116 | 0.548 | 6.794 | 0.509 | 9.852 | 0.516 |
| 250 | 3LQV_B | 12.904 | 0.575 | 9.519 | 0.511 | 11.862 | 0.613 | 10.770 | 0.433 |
| 251 | 3M1N_B | 18.313 | 0.437 | 16.966 | 0.385 | 16.851 | 0.492 | 18.227 | 0.351 |
| 252 | 3MQK_C | 2.229 | 0.737 | 3.363 | 0.559 | 2.227 | 0.754 | 9.480 | 0.370 |
| 253 | 3N1G_C | 3.659 | 0.711 | 4.063 | 0.611 | 2.616 | 0.818 | 6.229 | 0.450 |
| 254 | 3N9U_C | 6.612 | 0.585 | 6.560 | 0.576 | 6.730 | 0.758 | 8.843 | 0.584 |
| 255 | 3PD2_A | 7.310 | 0.569 | 7.753 | 0.541 | 4.189 | 0.723 | 9.737 | 0.467 |
| 256 | 3QU3_A | 7.832 | 0.478 | 7.147 | 0.452 | 5.686 | 0.508 | 10.597 | 0.379 |
| 257 | 3SDL_B | 25.206 | 0.307 | 24.890 | 0.310 | 12.272 | 0.295 | 26.655 | 0.280 |
| 258 | 3UE6_E | 7.889 | 0.550 | 7.337 | 0.576 | 5.437 | 0.797 | 10.843 | 0.414 |
| 259 | 3V1O_A | 11.452 | 0.414 | 8.348 | 0.413 | 4.816 | 0.662 | 10.556 | 0.351 |
| 260 | 3W1Z_D | 11.115 | 0.599 | 8.946 | 0.475 | 13.271 | 0.597 | 9.588 | 0.497 |
| 261 | 3X0G_A | 8.079 | 0.529 | 7.129 | 0.520 | 4.901 | 0.559 | 6.298 | 0.444 |
| 262 | 3X15_A | 14.241 | 0.491 | 10.413 | 0.394 | 19.236 | 0.445 | 15.231 | 0.493 |
| 263 | 4AIH_A | 2.531 | 0.799 | 2.809 | 0.763 | 5.952 | 0.643 | 2.790 | 0.766 |
| 264 | 4ASW_C | 2.932 | 0.712 | 3.729 | 0.573 | 2.699 | 0.710 | 4.650 | 0.513 |
| 265 | 4B0M_A | 8.732 | 0.459 | 6.483 | 0.516 | 7.960 | 0.510 | 7.565 | 0.415 |
| 266 | 4CXT_A | 6.338 | 0.577 | 6.162 | 0.573 | 9.878 | 0.706 | 8.595 | 0.508 |
| 267 | 4ESB_A | 2.086 | 0.825 | 3.077 | 0.720 | 4.888 | 0.687 | 3.261 | 0.729 |
| 268 | 4GDK_A | 5.310 | 0.569 | 4.442 | 0.542 | 2.739 | 0.696 | 5.440 | 0.518 |
| 269 | 4GF3_A | 7.533 | 0.582 | 8.229 | 0.545 | 5.577 | 0.658 | 8.478 | 0.436 |
| 270 | 4GQY_A | 8.401 | 0.587 | 10.859 | 0.360 | 6.807 | 0.742 | 8.373 | 0.505 |
| 271 | 4I60_A | 6.473 | 0.525 | 5.207 | 0.553 | 3.041 | 0.753 | 6.583 | 0.487 |
| 272 | 4IOS_A | 8.884 | 0.461 | 6.134 | 0.421 | 5.450 | 0.460 | 7.946 | 0.435 |
| 273 | 4J20_A | 5.960 | 0.583 | 4.331 | 0.584 | 4.336 | 0.553 | 12.666 | 0.377 |
| 274 | 4JGX_B | 4.624 | 0.698 | 4.922 | 0.671 | 4.669 | 0.647 | 6.276 | 0.488 |
| 275 | 4KA0_A | 4.856 | 0.643 | 4.260 | 0.609 | 2.319 | 0.849 | 5.477 | 0.536 |
| 276 | 4LE0_B | 1.901 | 0.849 | 2.476 | 0.776 | 1.361 | 0.914 | 2.334 | 0.821 |
| 277 | 4M75_F | 5.119 | 0.704 | 5.311 | 0.580 | 5.883 | 0.719 | 5.414 | 0.604 |
| 278 | 4MMG_A | 3.229 | 0.735 | 3.891 | 0.644 | 2.870 | 0.727 | 4.902 | 0.584 |
| 279 | 4OW1_A | 6.506 | 0.609 | 4.250 | 0.581 | 3.156 | 0.657 | 5.780 | 0.483 |
| 280 | 4Q2O_A | 5.457 | 0.644 | 4.124 | 0.656 | 6.629 | 0.776 | 8.061 | 0.546 |
| 281 | 4Q2Q_A | 4.134 | 0.632 | 5.285 | 0.591 | 7.642 | 0.703 | 6.483 | 0.622 |
| 282 | 4R67_0 | 10.917 | 0.344 | 15.811 | 0.254 | 2.406 | 0.864 | 15.196 | 0.416 |
| 283 | 4RUV_A | 2.334 | 0.786 | 2.510 | 0.742 | 1.788 | 0.850 | 2.208 | 0.822 |
| 284 | 4UIJ_A | 4.714 | 0.713 | 3.421 | 0.709 | 2.689 | 0.766 | 3.318 | 0.750 |
| 285 | 4V2O_A | 1.785 | 0.807 | 4.104 | 0.572 | 6.903 | 0.545 | 4.187 | 0.523 |
| 286 | 5CJ3_B | 5.147 | 0.603 | 3.786 | 0.708 | 5.503 | 0.627 | 8.510 | 0.511 |
| 287 | 5E4E_A | 15.330 | 0.308 | 15.702 | 0.248 | 4.518 | 0.610 | 14.948 | 0.256 |
| 288 | 5EKT_A | 12.904 | 0.439 | 9.065 | 0.415 | 5.261 | 0.690 | 10.720 | 0.350 |
| 289 | 5IAO_A | 11.531 | 0.467 | 10.414 | 0.426 | 4.440 | 0.690 | 14.361 | 0.326 |
| 290 | 5IZB_A | 11.344 | 0.458 | 10.870 | 0.460 | 14.473 | 0.557 | 13.162 | 0.422 |
| 291 | 5JTM_A | 9.969 | 0.455 | 10.140 | 0.439 | 11.441 | 0.580 | 12.437 | 0.284 |
| 292 | 5L38_A | 2.898 | 0.712 | 10.379 | 0.693 | 1.721 | 0.856 | 8.748 | 0.373 |
| 293 | 5L8R_D | 14.062 | 0.400 | 15.955 | 0.352 | 17.477 | 0.360 | 17.650 | 0.322 |
| 294 | 5O2V_A | 3.477 | 0.730 | 2.986 | 0.703 | 4.862 | 0.775 | 4.451 | 0.540 |
| 295 | 5O8G_A | 6.731 | 0.564 | 6.896 | 0.524 | 5.015 | 0.772 | 12.582 | 0.320 |
| 296 | 5T17_A | 2.799 | 0.717 | 2.893 | 0.657 | 2.634 | 0.724 | 3.213 | 0.659 |
| 297 | 5TMF_E | 9.924 | 0.523 | 7.720 | 0.507 | 5.786 | 0.505 | 11.066 | 0.563 |
| 298 | 5TUV_B | 13.091 | 0.435 | 12.824 | 0.411 | 12.283 | 0.388 | 14.670 | 0.432 |

Continued on next page

Continue to last page

| No. | Protein | SNfold | | Rosetta-dist(500) | | C-QUARK | | SNfold-exploitation | |
|---|---|---|---|---|---|---|---|---|---|
| | | RMSD (Å) | TM-score | RMSD (Å) | TM-score | RMSD (Å) | TM-score | RMSD (Å) | TM-score |
| 299 | 5WSE_A | 4.262 | 0.672 | 4.899 | 0.621 | 5.817 | 0.667 | 7.267 | 0.562 |
| 300 | 6AQ3_B | 7.493 | 0.523 | 6.989 | 0.470 | 11.971 | 0.508 | 10.971 | 0.344 |

Table S5: Results of SNfold and Rosetta-dist(1000) on 50 proteins.

| No. | Protein | SNfold | | | Rosetta-dist(1000) | | |
|---|---|---|---|---|---|---|---|
| | | RMSD (Å) | TM-score | FE | RMSD (Å) | TM-score | FE |
| 1 | 1A6L_A | 4.616 | 0.579 | 1.05E+06 | 3.942 | 0.579 | 1.97E+08 |
| 2 | 1A7D_A | 3.195 | 0.764 | 1.06E+06 | 4.097 | 0.733 | 1.83E+08 |
| 3 | 1AKP_A | 9.506 | 0.401 | 9.02E+05 | 9.236 | 0.433 | 1.80E+08 |
| 4 | 1BJX_A | 7.551 | 0.640 | 1.43E+06 | 7.094 | 0.661 | 2.69E+08 |
| 5 | 1CTO_A | 8.696 | 0.371 | 9.76E+05 | 5.572 | 0.506 | 1.83E+08 |
| 6 | 1ELW_A | 1.252 | 0.920 | 9.83E+05 | 1.383 | 0.906 | 1.48E+08 |
| 7 | 1F9P_A | 7.281 | 0.616 | 1.71E+06 | 5.749 | 0.575 | 3.38E+08 |
| 8 | 1FAQ_A | 3.476 | 0.551 | 9.53E+05 | 4.284 | 0.497 | 1.71E+08 |
| 9 | 1FCA_A | 1.184 | 0.821 | 9.64E+05 | 1.852 | 0.697 | 1.60E+08 |
| 10 | 1GPQ_B | 7.349 | 0.545 | 1.08E+06 | 6.610 | 0.598 | 1.63E+08 |
| 11 | 1I35_A | 4.816 | 0.603 | 9.73E+05 | 5.068 | 0.562 | 1.68E+08 |
| 12 | 1IUY_A | 8.803 | 0.621 | 9.99E+05 | 9.435 | 0.538 | 1.55E+08 |
| 13 | 1K73_1 | 3.853 | 0.670 | 8.61E+05 | 2.853 | 0.651 | 1.67E+08 |
| 14 | 1KN6_A | 3.864 | 0.586 | 9.79E+05 | 3.394 | 0.573 | 1.77E+08 |
| 15 | 1KSX_A | 5.780 | 0.540 | 9.50E+05 | 4.507 | 0.617 | 1.72E+08 |
| 16 | 1L1D_B | 8.146 | 0.376 | 1.02E+06 | 5.518 | 0.533 | 1.68E+08 |
| 17 | 1L2P_A | 0.622 | 0.949 | 8.06E+05 | 0.897 | 0.907 | 1.41E+08 |
| 18 | 1LR1_B | 3.867 | 0.536 | 9.44E+05 | 4.102 | 0.514 | 1.48E+08 |
| 19 | 1LZW_B | 4.827 | 0.752 | 1.05E+06 | 5.825 | 0.755 | 1.81E+08 |
| 20 | 1MFQ_C | 8.425 | 0.627 | 9.70E+05 | 8.905 | 0.602 | 1.57E+08 |
| 21 | 1N12_A | 11.512 | 0.353 | 1.06E+06 | 6.455 | 0.444 | 1.84E+08 |
| 22 | 1NOE_A | 7.189 | 0.392 | 1.17E+06 | 5.106 | 0.458 | 1.96E+08 |
| 23 | 1OFT_A | 3.309 | 0.702 | 9.66E+05 | 3.282 | 0.666 | 1.85E+08 |
| 24 | 1OZ9_A | 4.839 | 0.649 | 1.38E+06 | 7.080 | 0.608 | 2.52E+08 |
| 25 | 1PIH_A | 8.365 | 0.428 | 1.30E+06 | 7.053 | 0.415 | 2.21E+08 |
| 26 | 1PMS_A | 8.434 | 0.510 | 9.65E+05 | 12.034 | 0.553 | 1.64E+08 |
| 27 | 1TUL_A | 9.036 | 0.400 | 9.31E+05 | 10.256 | 0.252 | 1.83E+08 |
| 28 | 1TYG_B | 2.141 | 0.733 | 9.24E+05 | 2.228 | 0.734 | 1.76E+08 |
| 29 | 1UFB_A | 4.626 | 0.732 | 1.02E+06 | 3.817 | 0.746 | 1.72E+08 |
| 30 | 2ACY_A | 3.470 | 0.670 | 9.34E+05 | 3.315 | 0.668 | 1.87E+08 |
| 31 | 2AQ0_A | 7.690 | 0.658 | 1.48E+06 | 7.429 | 0.648 | 2.79E+08 |
| 32 | 2BSE_A | 10.783 | 0.441 | 8.90E+05 | 8.132 | 0.419 | 1.81E+08 |
| 33 | 2BYK_D | 1.764 | 0.870 | 9.72E+05 | 1.799 | 0.858 | 1.50E+08 |
| 34 | 2CDP_A | 10.584 | 0.371 | 9.89E+05 | 6.330 | 0.484 | 1.84E+08 |
| 35 | 2CMX_A | 2.404 | 0.719 | 1.00E+06 | 2.499 | 0.691 | 1.56E+08 |
| 36 | 2EWC_B | 5.272 | 0.639 | 1.10E+06 | 7.503 | 0.658 | 1.86E+08 |
| 37 | 2H30_A | 8.887 | 0.574 | 1.17E+06 | 6.492 | 0.643 | 1.80E+08 |
| 38 | 2NAZ_A | 4.904 | 0.670 | 9.36E+05 | 3.986 | 0.668 | 1.75E+08 |
| 39 | 2P7L_A | 4.850 | 0.627 | 7.85E+05 | 4.251 | 0.661 | 1.79E+08 |
| 40 | 2QVG_A | 3.535 | 0.752 | 9.21E+05 | 3.395 | 0.746 | 1.70E+08 |
| 41 | 3EOD_A | 4.011 | 0.795 | 7.31E+05 | 4.044 | 0.776 | 1.59E+08 |
| 42 | 3MQK_C | 2.229 | 0.737 | 9.01E+05 | 4.116 | 0.463 | 1.75E+08 |
| 43 | 3PD2_A | 7.310 | 0.569 | 1.01E+06 | 6.793 | 0.631 | 1.86E+08 |
| 44 | 3W1Z_D | 11.115 | 0.599 | 1.65E+06 | 7.734 | 0.558 | 3.07E+08 |
| 45 | 4ESB_A | 2.086 | 0.825 | 9.79E+05 | 3.032 | 0.712 | 1.55E+08 |
| 46 | 4JGX_B | 4.624 | 0.698 | 1.02E+06 | 4.643 | 0.696 | 1.69E+08 |
| 47 | 5E4E_A | 15.330 | 0.308 | 9.69E+05 | 15.127 | 0.219 | 1.68E+08 |
| 48 | 5L8R_D | 14.062 | 0.400 | 1.63E+06 | 14.029 | 0.336 | 3.51E+08 |
| 49 | 5O2V_A | 3.477 | 0.730 | 1.54E+06 | 2.750 | 0.740 | 2.82E+08 |
| 50 | 5WSE_A | 4.262 | 0.672 | 9.27E+05 | 4.636 | 0.649 | 1.89E+08 |

Table S6: Results of SNfold, QUARK, BAKER-ROSETTASERVER, RaptorX-DeepModeller and MULTICOM_CLUSTER on 24 FM targets from CASP13.

| Target | SNfold | QUARK | BAKER-ROSETTASERVER | RaptorX-DeepModeller | MULTICOM_CLUSTER |
|---|---|---|---|---|---|
| T0950-D1 | 0.51 | 0.44 | 0.46 | 0.56 | 0.22 |
| T0953s1-D1 | 0.45 | 0.40 | 0.19 | 0.28 | 0.38 |
| T0953s2-D1 | 0.31 | 0.36 | 0.33 | 0.18 | 0.20 |
| T0953s2-D2 | 0.39 | 0.48 | 0.47 | 0.69 | 0.22 |
| T0953s2-D3 | 0.15 | 0.35 | 0.19 | 0.29 | 0.14 |
| T0955-D1 | 0.53 | 0.73 | NA | 0.59 | 0.77 |
| T0957s1-D1 | 0.48 | 0.40 | 0.42 | 0.37 | 0.31 |
| T0957s2-D1 | 0.67 | 0.53 | 0.48 | 0.65 | 0.51 |
| T0958-D1 | 0.71 | 0.55 | 0.53 | 0.66 | 0.55 |
| T0960-D2 | 0.43 | 0.44 | 0.28 | 0.49 | 0.38 |
| T0963-D2 | 0.42 | 0.46 | 0.36 | 0.48 | 0.28 |
| T0968s1-D1 | 0.56 | 0.56 | 0.74 | 0.62 | 0.43 |
| T0968s2-D1 | 0.62 | 0.65 | 0.66 | 0.59 | 0.42 |
| T0969-D1 | 0.34 | 0.64 | 0.49 | 0.65 | 0.44 |
| T0970-D1 | 0.37 | 0.51 | 0.40 | 0.54 | 0.33 |
| T0980s1-D1 | 0.46 | 0.54 | 0.41 | 0.37 | 0.25 |
| T0990-D1 | 0.62 | 0.58 | 0.37 | 0.40 | 0.36 |
| T0990-D2 | 0.46 | 0.37 | 0.26 | 0.35 | 0.25 |
| T0990-D3 | 0.33 | 0.22 | 0.23 | 0.22 | 0.24 |
| T1005-D1 | 0.42 | 0.70 | 0.70 | 0.70 | 0.69 |
| T1008-D1 | 0.67 | 0.35 | 0.56 | 0.28 | 0.38 |
| T1021s3-D1 | 0.37 | 0.64 | 0.50 | 0.66 | 0.50 |
| T1021s3-D2 | 0.40 | 0.45 | 0.19 | 0.59 | 0.27 |
| T1022s1-D1 | 0.39 | 0.55 | 0.40 | 0.58 | 0.38 |
| Average | 0.461 | 0.496 | 0.418 | 0.491 | 0.371 |
